# Structure of Pex8 in complex with peroxisomal receptor Pex5 reveals its essential role in peroxisomal cargo translocation

**DOI:** 10.1101/2025.08.30.673231

**Authors:** Lakhan Ekal, Daniel Wendscheck, Yotam David, Grzegorz Chojnowski, Cy M. Jeffries, Edukondalu Mullapudi, Maya Schuldiner, Bettina Warscheid, Einat Zalckvar, Matthias Wilmanns

## Abstract

Peroxisomes are essential cellular organelles that enable the sequestered execution of a broad range of metabolic processes. Due to the lack of an internal protein synthesis machinery, they entirely depend on the import of target proteins to carry out their functions within peroxisomes. While the process of cargo/receptor recognition is well understood, knowledge about the molecular mechanisms of the subsequent translocation steps, including cargo release and receptor recycling, is lacking behind. Here, we provide structural and functional evidence on the role of Pex8 in these processes. First, we show that Pex8 in yeast is essential for peroxisomal cargo translocation, irrespective of the mechanism of receptor/cargo recognition. Next, we reveal that Pex8 binds through an irregular twelvefold HEAT repeat array to a short three-helical bundle within the otherwise unfolded N-terminal domain of the Pex5 receptor. Impairing this interaction abolishes peroxisomal protein translocation. It is complemented by a secondary autonomous Pex8 cargo-like interaction site with the C-terminal domain of Pex5, thus generating a bipartite interaction between the two proteins.

Our data support a model in which Pex5/Pex8 complex formation allows assembly with the peroxisomal Pex2/Pex10/Pex12 E3-ubiquitin ligase complex to initiate recycling of the receptor. In summary, our findings provide in-depth insight into the transition from cargo release into peroxisomes to receptor recycling, which is essential to uncover the overall process of the peroxisomal cargo translocation.

## Introduction

Peroxisomes are ubiquitous eukaryotic organelles that are responsible for multiple metabolic processes requiring sequestration, including the breakdown of fatty acids, and play major roles in human health and disease **[1, 2].** Compromised functions of peroxisomes due to the failure of proper protein import can lead to severe peroxisomal biogenesis disorders (PBDs) **[3, 4].** Over one hundred folded proteins that mostly harbor specific peroxisomal targeting signals (PTS) 1 or 2 are recognized by the peroxisomal import receptors Pex5 and Pex7, respectively **[2, 5, 6].** Upon cargo loading, these receptors dock and integrate within the docking/translocation machinery (DTM) complex, which minimally includes Pex13 and Pex14, and subsequently translocate the cargo across the peroxisomal membrane **[7, 8].** The DTM complex was proposed to form the basis of an inducible transient pore in the peroxisomal membrane upon cargo recognition **[9].** An alternative model has recently been suggested, in which a constitutive pore is formed for the translocation of cargo-loaded receptors into the peroxisomal lumen **[10–12].**

The aim of this study was to reveal how the primary peroxisomal import receptor Pex5 is regulated during the complex cargo translocation process, thereby promoting cargo release into the peroxisomal lumen and its subsequent recycling. At the molecular level, Pex5 is divided into a mostly unfolded N-terminal domain (NTD) and a C-terminal folded domain (CTD) **[13].** The CTD forms a sevenfold array of tetratricopeptide repeats (TPR), followed by a C-terminal three-helical bundle domain (CHB) **[5]**. The TPR array provides a highly conserved recognition site for peroxisomal cargoes carrying a C-terminal PTS1 motif. The Pex5 CTD also comprises auxiliary binding sites for some PTS1 cargoes with an additional recognition motif, referred to as PTS3, through a mostly helical extension toward the NTD **[14, 15].**

In contrast to the CTD, little is known about the mostly disordered structure of the Pex5 NTD **[13]**. In functional terms, the NTD is responsible for binding to the membrane component Pex13 and the docking factor Pex14 through short linear WxxxF and related sequence motifs **[5, 11, 16, 17]**. In addition, at the very N-terminus, Pex5 harbors an invariant cysteine, which is monoubiquitinated by the Pex2/Pex10/Pex12 Really Interesting New Gene (RING) finger complex with E3-ubiquitin ligase activity, as the initial step of receptor recycling **[18, 19]**.

Recent data indicate that a loose array of predicted helices near the Pex5 N-terminus is involved in docking with the E3-ubiquitin ligase complex **[10, 16]**. Various lines of evidence have shown that these processes depend on cargo loading primarily to the CTD of the receptor, but any mechanistic insight on how this signal is transmitted to the receptor NTD has remained elusive **[7, 20].**

In fungi and in plants, an additional protein factor Pex8 is implicated in this process **[21–23].** Although Pex8 sequences do not reveal any features typical of transmembrane proteins, Pex8 has consistently been detected to be associated with the peroxisomal membrane, suggesting that it interacts with components of the membrane-associated protein translocation machinery **[24]**. These findings are in agreement with Pex8 acting as a linker between the DTM and the E3-ubiquitin ligase complex **[20, 23, 25].** Unlike most other Pex5 PTS1 cargoes, Pex8 interacts with both the NTD and CTD of Pex5 **[26].** Interestingly, the presence of the PTS1 motif in Pex8 is dispensable for importing matrix proteins, especially for yeast cell growth on oleate-containing media where peroxisomes are vital for cell survival **[26–28].** Pex8 has also been reported to interact with the Pex7 coreceptor Pex20 via a putative PTS2 motif **[28]**. However, this site is not conserved in fungal and plant Pex8 sequences and its functional significance still remains unclear **[22].**

To elucidate the mechanistic role of Pex8 in peroxisomal cargo translocation, we investigated its structure and function in the absence and presence of the Pex5 receptor. First, we revealed an essential role for Pex8, independent of its PTS1 motif, in importing diverse cargoes. Using an integrative structural biology approach, we found that Pex8 folds into an irregular three-segmented array of 12 HEAT repeats with limited conformational flexibility between these segments. We discovered a bimodal interaction between Pex5 and Pex8, showing that Pex8 directly and independently binds to both the NTD, through the formation of a three-helical bundle within the otherwise unfolded NTD, and the CTD of Pex5. When the Pex5 NTD / Pex8 interaction was disrupted cargo import was impaired, demonstrating the essential functional role of this interaction. Our results support a model in which Pex8 acts as a structural adapter that facilitates the delivery of the Pex5 N-terminus for C10 mono-ubiquitination by the peroxisomal E3-ubiquitin ligase complex, thereby initiating the recycling process of Pex5. In summary, our study provides mechanistic insights into the role of Pex8 in cargo release, coupled with preparing the Pex5 receptor for recycling.

## Results

### Experiment Design

As a prerequisite for biophysical and structural analysis we screened Pex8 and Pex5 proteins from different yeast species on their suitability for *in vitro* functional and structural analysis. We purified an N-terminally truncated version of Pex8 (residues 33-713), full-length (FL) Pex5, and truncated versions of Pex5 that include either the NTD (residues 1-276) or the CTD (residues 259-576), all from *Pichia pastoris (P. pastoris)*. Functional experiments were carried out with the corresponding genes from *Saccharomyces cerevisiae (S. cerevisiae),* as the genetics of this organism are well established (**Figure 1a**). Pex5 and Pex8 sequences from *P. pastoris* and *S. cerevisiae* share 38% (57%) and 21% (36%) identity (similarity), respectively (**Supplementary Figures 1-2**). Structural predictions of Pex5 and Pex8 from both organisms suggest closely related folds of the respective protein components in complex with each other (**Supplementary Figures 3-4**). In particular, the well-established bipartite NTD/CTD domain organization of Pex5 is identical in both organisms. These close relationships provide the basis for the data presented to be valid for both yeast species.

**Figure 1.**
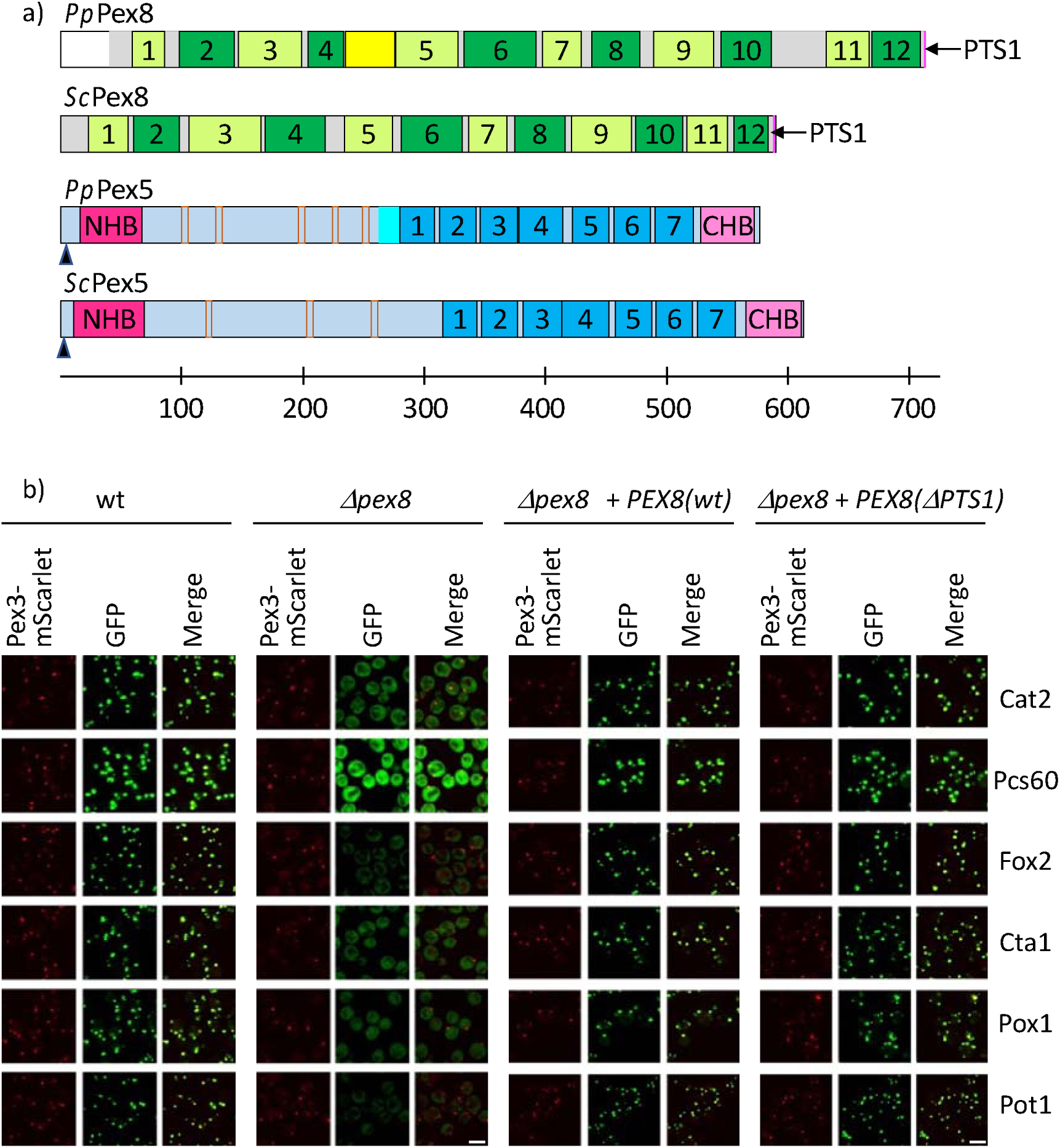
Pex8 is generally required for peroxisomal cargo import. **a**, scheme of the structural/functional organization of the related *P. pastoris (Pp)* and *S. cerevisiae (Sc)* Pex5 and Pex8 sequences (for further details on structural elements see Figures 4-5). Pex8 HEAT repeats are in alternating colors (limon, green) and are numbered 1-12. The structured insert-1 domain, connecting segments I and II, and the C-terminal PTS1 motif of Pex8 are colored in yellow and magenta, respectively. The Pex5 cysteine mono-ubiquitination site is indicated by a black vertical arrow. The NHB is in hot-pink, the degenerate WxxxF motifs are in brown, the extended folded segment of the Pex5 CTD is in cyan (only shown for *P. pastoris* Pex5, for which experimental structural evidence is available), the sevenfold repeated array of TPRs is in marine and numbered, the CHB is in violet. All functional and structural elements shown are proportional to the sequence scale bar below; **b**, fluorescence microscopy images showing co-localization of cargoes Cat2, Pcs60, Fox2, Cta1, Pox1 and Pot1 with peroxisomes (Pex3) in WT and *pex8*Δ *S. cerevisiae* cells, complemented with *PEX8* or *PEX8(*Δ*PTS1).* Scale bars: 5 µm.

### Pex8 is generally required for peroxisomal cargo import

To test whether there is a general role of Pex8 in peroxisomal cargo translocation, we selected six different cargoes that were shown to have different Pex5 modes of interaction **[29]**: four contain a C-terminal PTS1 motif (Cat2p, Fox2p, Cta1p, Pcs60), one harbors an N-terminal PTS2 motif (Pot1) and the last one has neither a PTS1 nor a PTS2 motif (Pox1) **[29].** Pox1 binding to the Pex5 receptor was assigned to a mostly helical extension of the Pex5 CTD **[30, 31].** For all selected PTS1 cargoes, additional Pex5 binding sites were reported, in part overlapping with the site in Pox1, also referred to as the PTS3 binding site **[14, 32]**. Unlike all other selected cargoes in this screen, Pex8 translocation on its own was sensitive to a synthetic PTS1 competitor **[29]**, indicating inferior binding properties to the Pex5 receptor.

Here, we first analyzed *PEX8*-deficient (*pex8*Δ*) S. cerevisiae* strains, each expressing one of the six selected N-terminally GFP-tagged cargoes, by epifluorescence microscopy (**Figure 1b).** None of these strains exhibited proper peroxisomal cargo localization in the absence of Pex8, demonstrating a functional role of Pex8 in cargo translocation to be general and independent from the presence of any specific receptor recognition motif. When these strains were complemented with plasmids encoding either wild-type (WT) *PEX8* or *PEX8* lacking the C-terminal PTS1 motif (*PEX8(*Δ*PTS1)*), proper translocation of all cargoes could be restored. Therefore, the data neither indicated that the PTS1 motif of Pex8 plays a primary role in promoting proper translocation of other cargoes nor supported the existence of a Pex8-mediated direct competition mechanism against PTS1 cargo binding to the receptor.

Ultimately, the data were suggestive of the involvement of other PTS1-independent binding sites between the Pex5 receptor and Pex8 to carry out its function in peroxisomal cargo translocation.

### Pex8 autonomously interacts with the Pex5 NTD and CTD

To characterize the interaction between Pex5 and Pex8 quantitatively, we employed Biolayer Interferometry (BLI-Octet) (**Figure 2a-c, Supplementary Figure 5**). In these experiments, Pex8 showed a dissociation constant K_D_ of 78 nM toward Pex5 FL, which is within the range observed for several peroxisomal cargoes **[5, 29, 33].** The K_D_ of Pex8/Pex5 CTD interaction was only 4.4 μM, which is equivalent to a more than 50-fold reduction in binding affinity, compared to Pex5 FL. This contrasts with observations on various PTS1 cargoes, in which PTS1-mediated interactions dominate over other secondary cargo/receptor binding sites **[32–34].** No binding was detected when using Pex5 CTD and Pex8 (ΔPTS1) constructs, confirming that the interaction between the Pex5 CTD and Pex8 is mediated by the C-terminal PTS1 motif of Pex8. Conversely, when either the PTS1 motif in Pex8 or the CTD of Pex5 was removed, abolishing PTS1-mediated Pex5 TPR binding, the dissociation constants were 478 nM and 683 nM, respectively, which is equivalent to a six-to eight-fold reduction in the strength of the Pex5/Pex8 interaction. The data thus demonstrate that despite the PTS1-mediated interaction being relatively weak on its own, it significantly contributes to the overall Pex5/Pex8 interaction.

**Figure 2.**
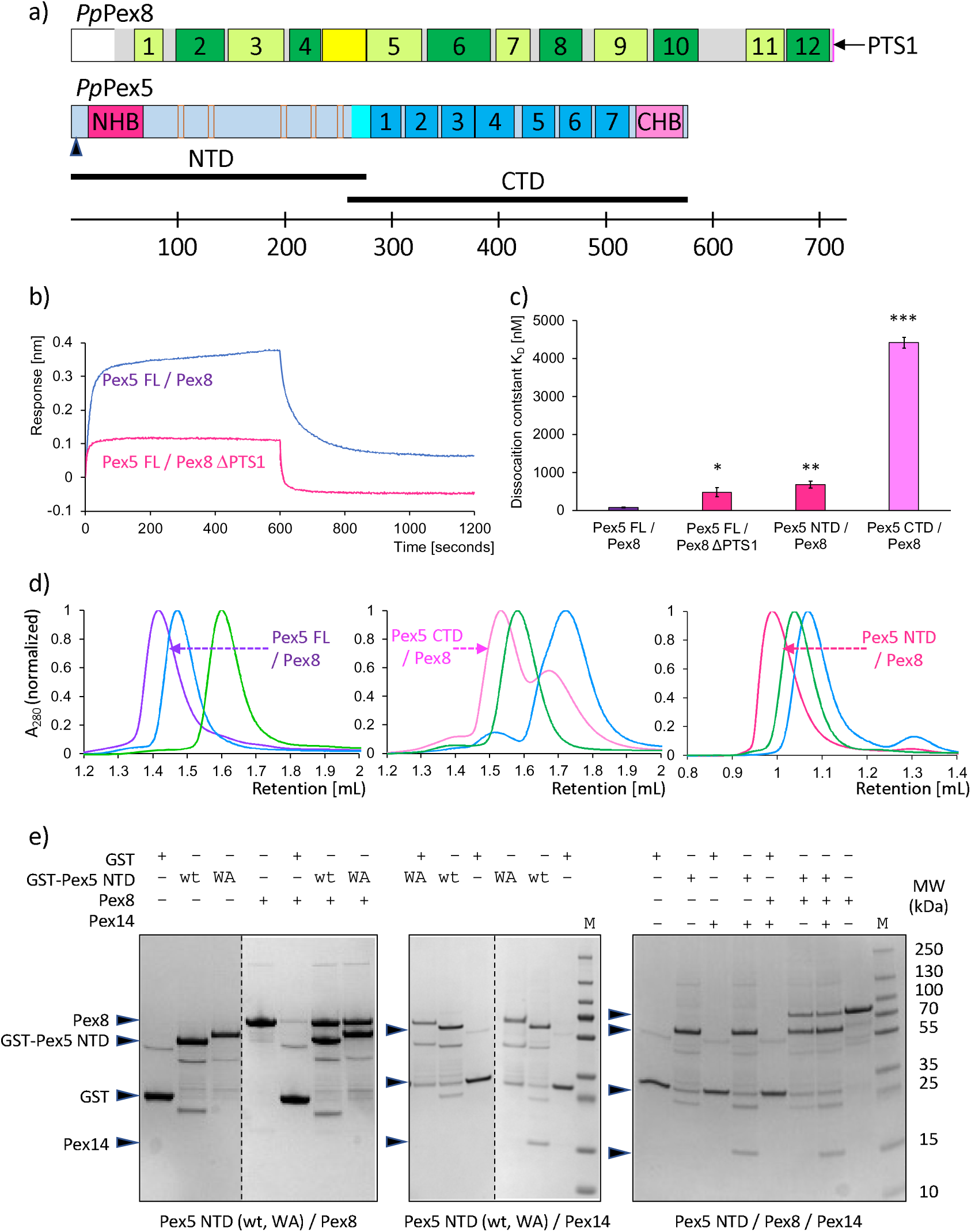
Pex8 interacts autonomously with the Pex5 NTD and CTD. **a**, scheme of the structural/functional organization of the *P. pastoris (Pp)* Pex5 and Pex8 sequences. Colors are as in Figure 1a. The background of the construct sequences used for characterization are in shaded in faint grey and blue. The extent of the Pex5 NTD and CTD constructs are indicated by additional black horizontal bars. **b,** representative BLI Octet data of Pex5 FL interacting with Pex8 WT and ΔPTS1, respectively (see also **Supplementary Figure 5**); **c,** dissociation constants K_D_ of different Pex5/Pex8 complexes. Color codes: both Pex5 NHB/Pex8 and Pex5 CTD/Pex8 PTS1 interaction sites intact, violet; only the Pex5 CTD / Pex8 PTS1 interaction site intact, magenta; only Pex5 NHB/Pex8 interaction site intact, hot-pink; The data are represented as the mean of N = 3 independent biological replicates ± SD. Statistical significance was determined using a one-tailed, unpaired Student’s t-test; p=0.00015 < 0.001 (***), p=0.0036 < 0.01 (**), p=0.0133 < 0.05 (*). **d**, Normalized size exclusion chromatography (SEC) profiles of different Pex5 constructs in complex with Pex8, color codes are as in panel c. The SEC profiles of the separate Pex5 and Pex8 constructs are in marine and green, respectively. Data were obtained from analytical S200 increase 3.2/300 columns (left and central panels) and S75 3.2/300 increase (right panel) columns. For further data, see **Supplementary Figure 7**. **e,** Pex5 NTD/Pex8/Pex14 interactions, based on GST-Pex5 NTD pulldown assays using Coomassie-stained SDS-PAGE gels: left panel, Pex5 NTD/Pex8 interactions are unaffected in Pex5 NTD variants with five mutated WxxxF (WA) motifs; middle panel, Pex14 interactions are affected by Pex5 NTD WA variant; right panel, Pex5 NTD can form a ternary complex with Pex14 and Pex8. Labeled marker lanes are shown to the right.

Our BLI measurements were supported by isothermal titration calorimetry (ITC) data, which confirmed a 1:1 binding stoichiometry and revealed that the interaction is enthalpy-driven, consistent with a strong network of specific contacts (**Supplementary Figure 6**). Stable complex formation between different Pex5 and Pex8 constructs, as a prerequisite for biophysical and structural analysis, was further shown by size exclusion chromatography data (**Figure 2d, Supplementary Figure 7**). In summary, our observations demonstrate that, in contrast to other PTS1 cargoes, bimodal binding between Pex8 and Pex5 is primarily mediated by a non-PTS1 interaction specifically involving the Pex5 NTD.

### Pex8 forms a non-canonical bimodal complex with Pex5 with limited conformational heterogeneity

To structurally study the impact of the bimodal Pex5/Pex8 interaction on peroxisomal import, we first used Small Angle X-ray Scattering (SAXS). Shape analysis of Pex5 FL and Pex8 on their own revealed elongated particles of both components, as expected for proteins with repeated helical modules (**Figure 3a-c, Supplementary Figure 8, Supplementary Table 1a**). Whereas the Pex8 SAXS data could be well interpreted by a single model, the Pex5 FL SAXS data required an ensemble modeling approach to properly account for the expected conformational heterogeneity of the mostly intrinsically disordered Pex5 NTD. The resulting models did not indicate any distinct orientation of the Pex5 NTD with respect to Pex5 CTD, confirming the highly flexible nature of the Pex5 receptor in the absence of bound protein ligands **[35, 36].**

**Figure 3.**
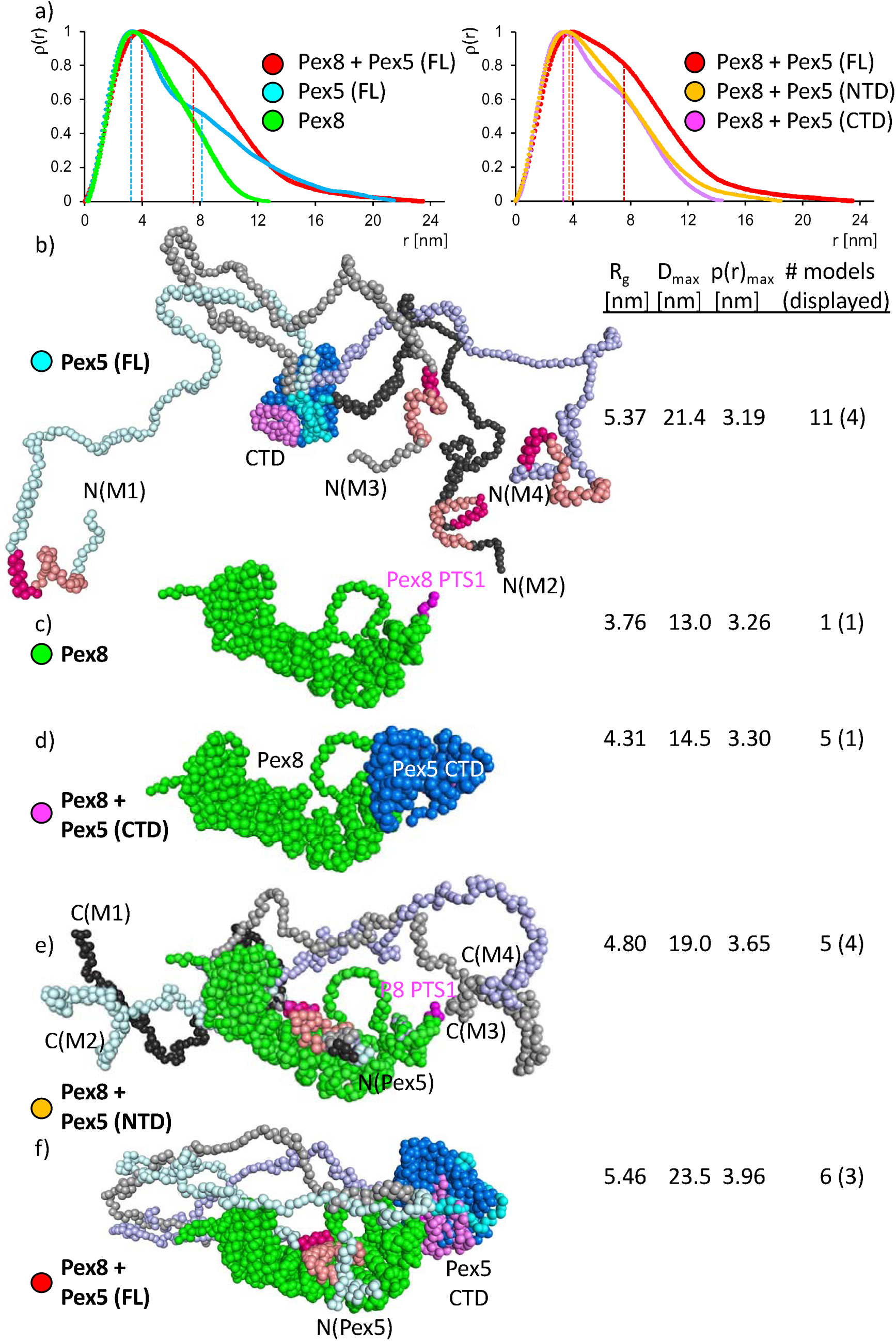
Pex8 forms a non-canonical bimodal complex with Pex5 with limited conformational heterogeneity. **a**, distance distribution p(r) plots of different Pex5/Pex8 complexes, calculated from SAXS curves (**Supplementary Figure 8**): left panel, Pex5 FL/Pex8 in comparison to separate Pex5 FL and Pex8; right panel, Pex5 FL/Pex8 complex in comparison to Pex5 NTD and CTD fragments in complex with Pex8. Ensemble models of separate Pex5 FL (**b**), single model of separate Pex8 (**c**), ensemble models of Pex5 CTD/Pex8 (**d**), Pex5 NTD/Pex8 (**e**), and Pex5 FL/Pex8 (**f**). For clarity only, subsets of ensemble models are shown (for illustration of all ensemble models, see **Supplementary Figure 8).** Models are shown in C_α_ sphere presentation. Color codes: Pex5 NHB, salmon (H1, H2) and hot pink (H3); remaining Pex5 NTD, different faint colors, depending on the number of ensemble models shown; Pex5 CTD, cyan (N-terminal extension), marine (TPR array) and violet (CHB); Pex5, green (HEAT repeats) and magenta (PTS1 motif). Some domains, motifs and termini are labeled, for guidance. For further details of structural elements, see Figures 4-5. Key SAXS parameters R_g_, D_max_, p(r)_max_, and the number of models required to obtain reasonable fits are listed for each model ensemble. The number of models shown is in parentheses. For further details, see **Supplementary Table 1.**

When complexed with different Pex5 versions (FL, NTD, and CTD), Pex8 consistently assembled in a 1:1 stoichiometry, in agreement with our ITC data (**Figure 3a, Supplementary Figure 6, Supplementary Table 1b**). To extract shape information of these complexes fitting the experimental SAXS data, initial structural models were generated based on AlphaFold3 (AF3) predictions and refined (**Supplementary Figure 3**). High-probability contacts in these models revealed two intermolecular interaction clusters, one corresponding to the expected Pex5 CTD/PTS1-mediated Pex8 site and the other one involving a previously uncharacterized interaction between the Pex5 NTD and the central part of Pex8 (**Supplementary Figure 4, Supplementary Table 2a**). Cohorts of 5-6 ensemble models were sufficient to provide robust fits with the experimental SAXS curves (**Figure 3d-f, Supplementary Figure 8).**

In these models, the Pex5 CTD interacting with the PTS1 motif of Pex8 consistently adopted closely related conformations, independent of the presence or absence of the Pex5 NTD (**Figure 3d, f**). Conversely, in all complexes with the Pex5 NTD present, only a short Pex5 segment covering residues 45-67 interacting with Pex8 was preserved (**Figure 3 e-f**). A preferred orientation of the remaining intrinsically disordered NTD (residues 68-260) along Pex8 could only be detected for Pex8 interacting with Pex5 FL, due to the second Pex5 CTD/Pex8 PTS1 interface. Taken together, our SAXS data suggest that both interfaces of Pex8 with the Pex5 NTD and Pex5 CTD are required for Pex5 to adopt a defined overall arrangement that parallels the elongated particle shape of Pex8. Due to the largely unstructured Pex5 NTD, the corresponding arrangements remain heterogenous and could only be modeled as ensembles.

### Pex8 interacts through a segmented HEAT repeat structure with both the Pex5 NTD and CTD

To decipher the structure of the Pex5 FL/Pex8 complex at high resolution, we used single-particle cryo-EM (**Figure 4, Supplementary Figure 9, Supplementary Table 3**). Initial 2D classification analysis showed that the complex particles are approximately 150 Å in length and consist of an elongated curved shape with well-defined features in the central section and more flexible regions with diffuse density at the two opposite poles (**Figure 4b**). Subsequent 3D *ab initio* reconstruction and refinement analysis resulted in a density map at an overall 4.5 Å resolution (**Figure 4c**). An atomic structural Pex5/Pex8 model was built into this density, by using the ensemble of SAXS-refined AF3 models, combined with *ab initio* modeling where feasible (**Figure 4d-e, Supplementary Figure 8e**). The resulting model was supported by experimental density for most of the Pex8 sequence including the C-terminal PTS1 motif, except for an unstructured N-terminal segment (residues 33-42) and a disordered insert close to its C-terminus (residues 592-628) (**Figure 4a,e**). For Pex5, the entire CTD (residues 264-576) and a well separated short helical fragment of the Pex5 NTD could be located in the resulting density (for further details see below). Different from most available high-resolutions structures of the Pex5 CTD **[5]**, in this structure also a preceding sequence segment (residues 264-274) was visible, which defines the boundary to the mostly unstructured Pex5 NTD. This segment has multiple interactions with the first two TPR modules and the CHB of the Pex5 CTD (**Figures 4e and 5d**).

**Figure 4.**
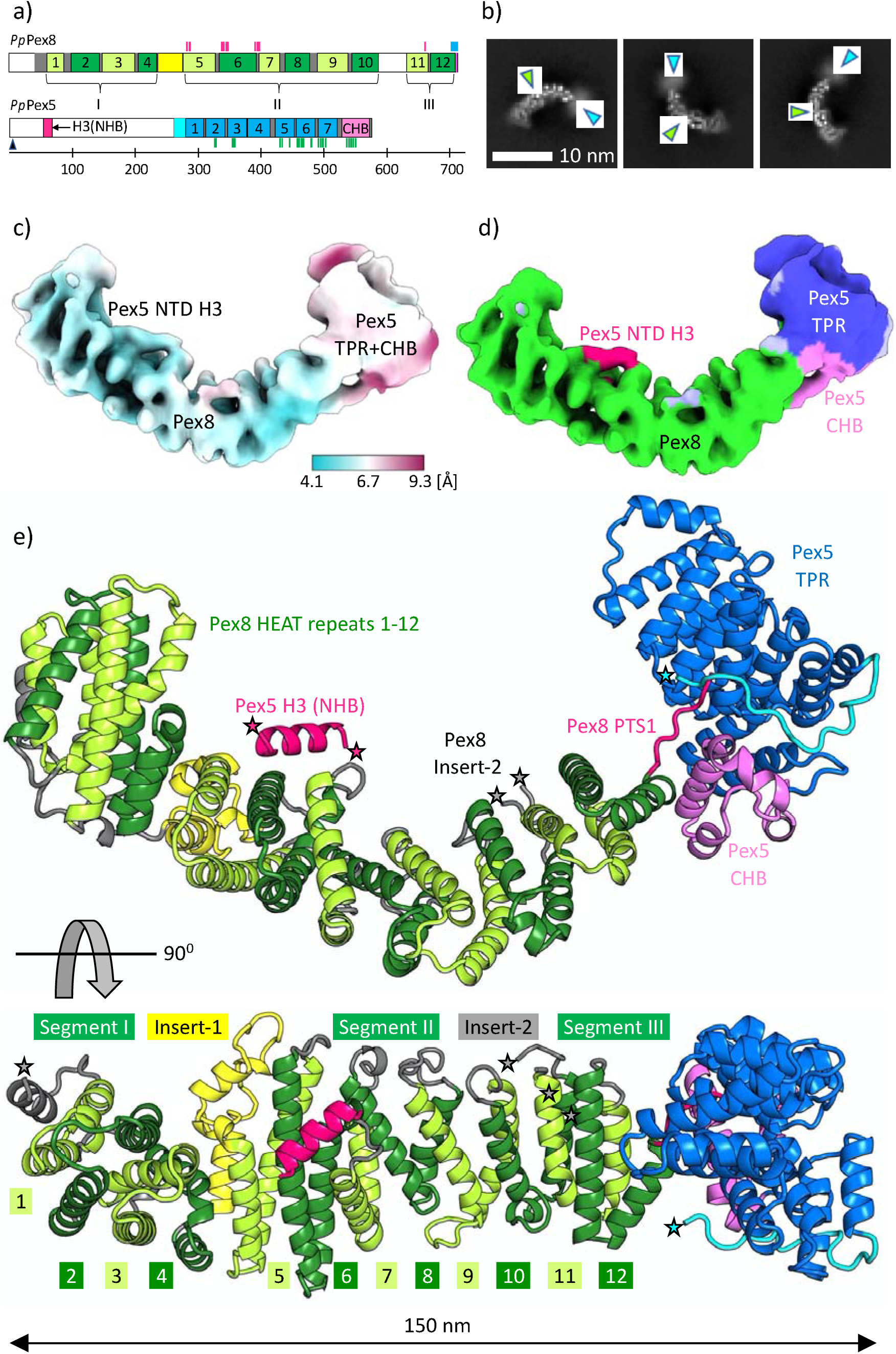
Pex8 interacts through a segmented HEAT repeat structure with both the Pex5 NTD and CTD. **a**, scheme of the structural/functional organization of the *P. pastoris (Pp)* Pex5 and Pex8 sequences. Colors are as in Figure 1a, except for sequence segments that remained invisible in the cryo-EM structure are left uncolored. Pex5/Pex8 interaction sites determined by cryo-EM analysis are indicated by vertical bars in complementary colors of the interaction partner; **b,** representative 2D class averages from the cryo-EM analysis, with rigid and flexible regions indicated by green and cyan arrows, representing the Pex5 CTD and Pex8, respectively; **c**, cryo-EM density map colored by local resolution, as indicated by the color range bar below, and **d,** applying a segmented coloring scheme, representing Pex8 (green), the Pex5 NHB H3 (hot pink, for further details see Figure 5), the Pex5 CTD TRP array (marine) and the Pex5 CTD CHB (violet). The remaining density that was assigned to Pex5 is colored in faint blue; **e**, cartoon representation of the cryo-EM Pex5/Pex8 complex structure in two different orientations rotated by 90 deg. around a horizontal axis, using the established coloring scheme (*cf*. panel a). The Pex8 HEAT repeats are numbered. For guidance, some of the structural/functional elements of Pex5 and Pex8 are labeled. Asterisks in colors matching the sequence fragments show where the segments are missing in the cryo-EM structure.

**Figure 5.**
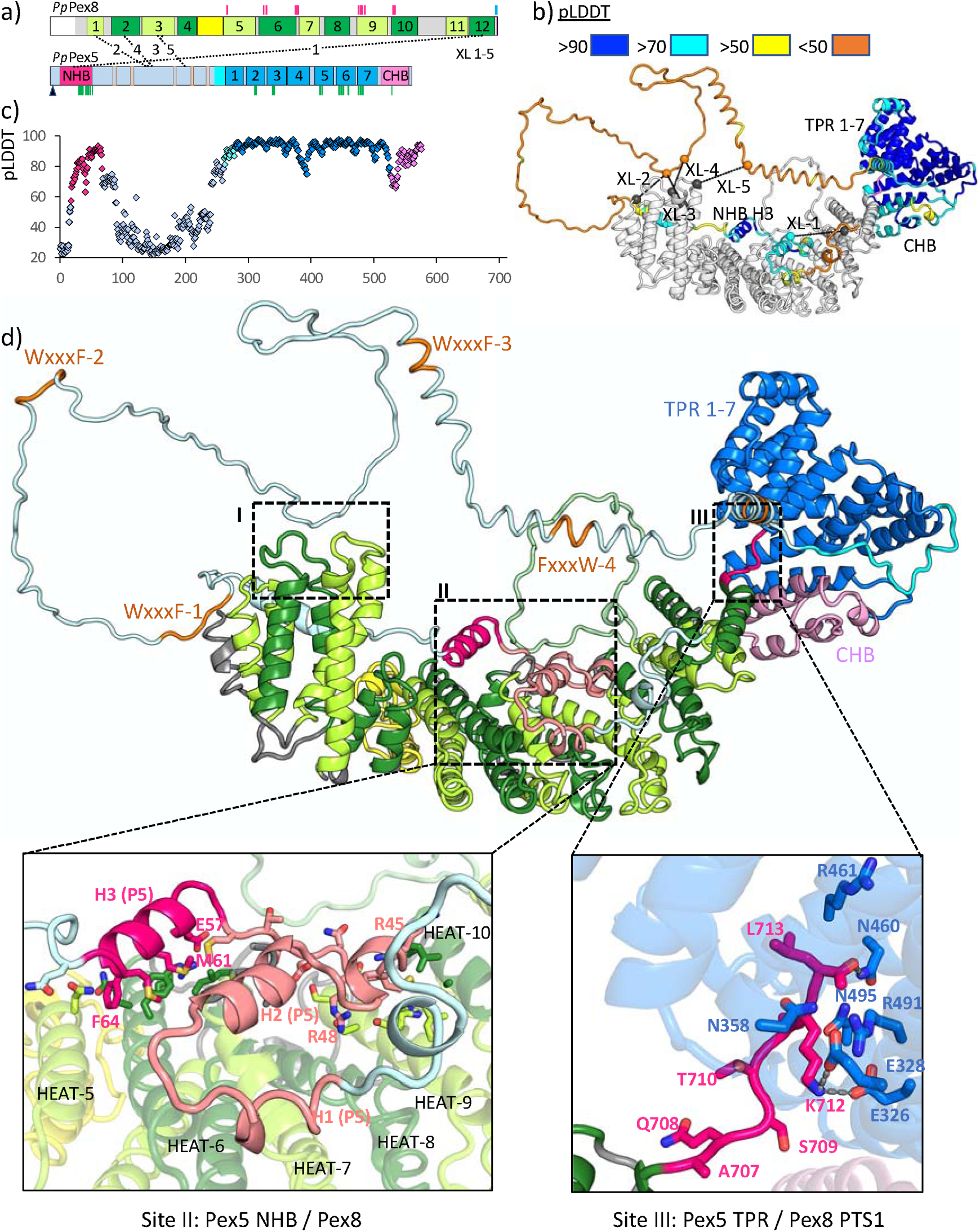
Structural characterization of the bimodal Pex5 NTD/Pex8 and Pex5 CTD/PTS1-mediated Pex8 interaction sites. **a**, scheme of the structural/functional organization of the *P. pastoris* Pex5 and Pex8 sequences. Colors are as in Figure 1a. Pex5/Pex8 interaction sites determined by cryo-EM analysis (*cf*. Figure 4a) and integrative modeling are indicated by vertical bars in complementary colors of the interaction partner. Intermolecular Pex5/Pex8 crosslinks are indicated by dashed lines (for further details see **Supplementary Figure 12, Supplementary Table 6) b,** experimentally restrained representative integrative Pex5/Pex8 model, showing Pex5 in AF3 pLDDT coloring scheme **[49].** The five intermolecular Pex5/Pex8 crosslinks (*cf*. panel a) are indicated. Some structural/functional elements are labeled for guidance. The complete model ensembles used for characterization of the Pex5/Pex8 interaction sites are shown in **Supplementary Figures 13-14. c,** Pex5 pLDDT residue scores, extracted from the representative model shown in panel b, indicating a high-confidence values for the folded CHB segment of the Pex5 NTD. Color codes are adapted from panel a. **d,** representative integrative Pex5/Pex8 model (same as shown in panel b), using color codes of panel a. Inlets II and III zoom in on structural details of the two Pex5 NHB/Pex8 and Pex5 CTD/Pex8 PTS1 interaction sites. Some structural/functional elements and residues of the two interaction sites, including all that were mutated for further biophysical and functional characterization of the Pex5 NHB/Pex8 interaction site (*cf*. Figure 6), are labeled for guidance. Potential hydrogen bonds between Pex5 CTD and Pex8 PTS1 residues of the respective interaction site are shown, where unambiguously predicted by homology modeling from known Pex5 CTD/PTS1 cargo high-resolution structures **[5, 6]**. Inlet I boxes an additional neighborhood site between Pex5 and Pex8 without evidence of specific residue/residue interactions, which was established by structural refinement of the intermolecular crosslinking data (*cf*. panels a and c). For reasons of space, “P5” labels are used as acronyms of Pex5.

To provide insight into the structural details of Pex8 at high resolution, we also solved its structure by X-ray crystallography at a 2.4 Å resolution (**Supplementary Figure 10**). Similar to the cryo-EM density, the X-ray density allowed modeling of residues 43-584 and 629-706 of the Pex8 sequence (**Figure 4, Supplementary Figure 10, Supplementary Table 4**). However, the C-terminal tail of Pex8 (residues 707-713) harboring the PTS1 motif in the absence of the Pex5 receptor was invisible. In both structures, Pex8 folds into an array of 12 HEAT repeat modules that form an irregular α-solenoid shape with a concave inner helical surface and a convex outer helical surface. The inner surface curvature, measured by angles between neighboring helices, completes almost one turn (353 degrees in the X-ray structure, 345 degrees in the cryo-EM structure) (**Figure 4e, Supplementary Figure 11a, Supplementary Table 5).**

We also found that the Pex8 HEAT repeat array is interspersed by two major inserts of different nature. The first insert between HEAT repeats 4 and 5 (residues 236-280) is well-structured and comprises two additional helices that do not follow the HEAT repeat pattern. This insert generates a major kink of close to 90 degrees between HEAT repeats 4 and 5 (**Figure 4e, Supplementary Figures 10b and 11a**). In contrast, the second insert between HEAT repeats 10 and 11 (residues 585-628) is unstructured and does not cause significant kinking of the overall α-solenoid. These two inserts divide the overall HEAT repeat array of Pex8 into three segments: segment I (HEAT repeats 1-4), segment II (HEAT repeats 5-10), and segment III (HEAT repeats 11-12) (**Figure 4a,e, Supplementary Figure 10b).**

Comparison of the cryo-EM and crystal structures of Pex8 revealed substantial differences in the overall arrangement of the HEAT repeat array, with spatial deviations of the opposite poles well exceeding 10 Å distances (**Supplementary Figure 11c-e**). These deviations are caused by a cumulative effect of relatively minor changes in the arrangement of neighboring HEAT repeats, induced by binding to Pex5 and reflecting an overall intrinsic flexibility typical for modular helical repeat array structures **[37–39]**.

### Structural characterization of the bimodal Pex5 NTD/Pex8 and Pex5 CTD/PTS1-mediated Pex8 interaction sites

To obtain an experimentally validated integrative and complete Pex5/Pex8 model, we used crosslinking-mass spectrometry (XL-MS), applying Sulfo-NHS-Diazirine (Sulfo-SDA) as crosslinker. From four independent experiments, we obtained five crosslinks that were found in at least three replicates (**Figure 5a-b, Supplementary Figure 12a,c, Supplementary Table 6a**). All crosslinked peptides were exclusively from the Pex5 NTD and from the two N-terminal HEAT repeats 1 and 2 of Pex8, except one from the C-terminal HEAT repeat 12. Using these crosslinks combined with AlphaFold-Multimer modeling, the fragmented cryo-EM/SAXS supported Pex5 CTD/Pex8 structure was expanded into a Pex5 FL/Pex8 model ensemble, in which all crosslinks were within permissive distances below 25 Å (**Figure 5b,d, Supplementary Figures 13-14, Supplementary Table 6b).** The models revealed additional loose neighborhood interactions between the N-terminal pole of the Pex8 segment I HEAT repeat structure and an unstructured Pex5 NTD segment between the NHB Pex8 interaction site and the Pex5 CTD.

To investigate whether this transient interaction site persists in the absence of the additional Pex5 CTD/Pex8 PTS1 interaction, we repeated the XL-MS analysis with Pex8 (ΔPTS1) (**Supplementary Figure 12b,d, Supplementary Table 6a**). Since all crosslinks of this dataset were identical to those in the Pex5/Pex8 WT dataset, we concluded that removal of the Pex5 CTD / Pex8 PTS1 binding site did not affect this additional transient Pex5 NTD / Pex8 interaction site, indicating that the observed overall increase in conformational heterogeneity of the Pex5/Pex8 complex (**Figure 3e**) is caused by regions of both binding partners not contributing to complex formation.

We further used the SAXS/EM/XL-MS restrained Pex5/Pex8 model ensemble for in-depth structural characterization of the two constitutive Pex5 NHB and CTD interaction sites with Pex8. Since the resolution of the Pex5 CTD segment in the overall density of the cryo-EM structure was considerably lower than that of most of Pex8, it became evident that its position is wobbly, in agreement to our SAXS data characterizing this interaction, and was therefore best captured by an ensemble of related conformers (**Figure 4c, Supplementary Figures 8c,e and 13-14**). Despite the fuzziness of the overall interaction, all of our models support a common binding mode of the C-terminal PTS1 sequence AKL (residues 711-713) of Pex8 into the well-established binding groove of the ellipsoidal TPR array of the Pex5 CTD, supported by a highly conserved set of specific interactions observed in reported Pex5/cargo structures (**Figure 5d, inlet; Supplementary Table 2a**). Therefore, we concluded that the imprecise positioning of the Pex5 CTD on the C-terminal pole of Pex8 is most likely due to the lack of a defined secondary interface, unlike observations in most Pex5 receptor/PTS1 cargo complex structures **[5, 35].**

We also used the integrative Pex5/Pex8 model ensemble to further characterize the distinct helical density from our cryo-EM data, next to the inner surface helices of HEAT repeats 5 and 6, and the loop connecting HEAT repeats 6 and 7 (residues 391-396) (**Figure 4d-e, Supplementary Figure 15).** Several residues from these segments are highly conserved among Pex8 sequences (**Supplementary Figures 2 and 15c**). AF3 models of the full-length Pex5/Pex8 complex indicated this density at high-confidence levels to be presented by the third helix (residues 54-66) of the three-helical bundle in the Pex5 NTD, which we assigned as NTD H3 (**Figures 4e and 5d, Supplementary Figure 3a,c**). This helix comprises one of the only two invariant residues (F64) in all Pex5 NTD sequences (**Figure 5d, Supplementary Figure 2**), supporting a crucial role of this interaction. Our models also consistently indicated an extension of the NTD H3 binding site toward the Pex5 N-terminus, supported by a specific crosslink (XL-1) connecting this segment to Pex8 (**Figure 5a-b,d, Supplementary Figures 13a and 14a, Supplementary Table 6**). This sequence stretch includes two additional predicted helices that we annotated as H1 (residues 16-22) and H2 (residues 28-36), which together with their connecting loops bind to the most conserved inner surface helices of Pex8 HEAT repeats 7, 8, 9 and 10 (**Figure 5d, Supplementary Figure 15**). Especially for the loop between helices H2 and H3, various additional high probability interactions with Pex8 were predicted (**Supplementary Table 2a).** In the presence of Pex8, these three Pex5 NTD helices H1, H2 and H3 form a N-terminal helical bundle (NHB), which is highly conserved in all structural models that are restrained by the available SAXS, EM, and XL-MS data (**Figure 5d, Supplementary Figures 13a and 14a).** In AF3 predictions, this region shows a marked peak in high predicted Local Distance Difference Test (pLDDT) confidence levels, framed by low-confidence regions, and reveals the sole major Pex5/Pex8 interaction site with well-defined distances in complementary AF3 Predicted Aligned Error (PAE) plots (**Figure 5c, Supplementary Figure 3b).** Thus, our data support the presence of an additional folded and well-defined Pex5 NTD segment when bound to Pex8, unlike Pex5 on its own (**Figure 5b-d, Supplementary Figures 8d-e, 13a and 14a**). An elevated curvature of the inner helix arrangements, especially of HEAT repeats 6, 7 and 8 in the range of 50 degrees between neighboring helices, generates a groove-like shape on the inner Pex8 surface, which allows a snug fit of the Pex5 NHB (**Supplementary Figure 11a).** As Helix H1 is directly preceded by the highly conserved N15 next to the mono-ubiquitination C10 site, Pex8 binding to the Pex5 NHB may also facilitate the initiation of Pex5 recycling **[18].**

Taken together, our models indicate a specific mode of Pex5 NTD/Pex8 binding, where the structurally conserved and folded NHB segment of the Pex5 NTD interacts with the Pex8 HEAT repeat array sequence in an approximately anti-antiparallel orientation. Subsequently, a disordered Pex5 NTD segment, adjacent to the NHB region, reorients through loose neighborhood interactions with the N-terminal pole of the Pex8 HEAT repeat structure, as indicated by the set of intermolecular crosslinks (**Figure 5b,d**). The remaining Pex5 NTD sequence turns toward the folded Pex5 CTD structure that is bound to the C-terminal pole of Pex8 through its PTS1 motif.

### Interaction between the Pex5 NHB and Pex8 is essential for cargo translocation

To experimentally confirm a functional role of the Pex5 NHB interaction with Pex8, we mutated R45, R48, E57, M61 and the invariant F64 on the NHB helix H3 (**Figures 5d (left inlet) and 6a**), to either reverse the charge or eliminate the functional role of their side chains. In pulldown assays of corresponding Pex5 NTD variants, significant loss of binding was achieved by the Pex5 single E57R mutant and the double mutants R45E/R48E and M61A/F64A (**Figure 6b**). These effects were subsequently quantified by BLI-Octet measurements, indicating a six-to eightfold increase in dissociation constants when compared to the interaction with Pex5 NTD WT, which is comparable to the effects observed for versions of the Pex5/Pex8, in which the Pex5 CTD/Pex8 PTS1 interaction was abolished (**Figure 6c-d, Supplementary Figures 16-17**). These data thus demonstrate that the NHB helix 3 and the preceding loop are crucial for the interaction with Pex8. We also confirmed that binding of the Pex5 NTD to the DTM scaffold component Pex14 via degenerate WxxxF motifs remains unaffected in the presence of Pex8, which was expected as none of these motifs are involved in Pex5 NTD/Pex8 binding (**Figures 2e and 5a,d, Supplementary Figure 1**).

**Figure 6.**
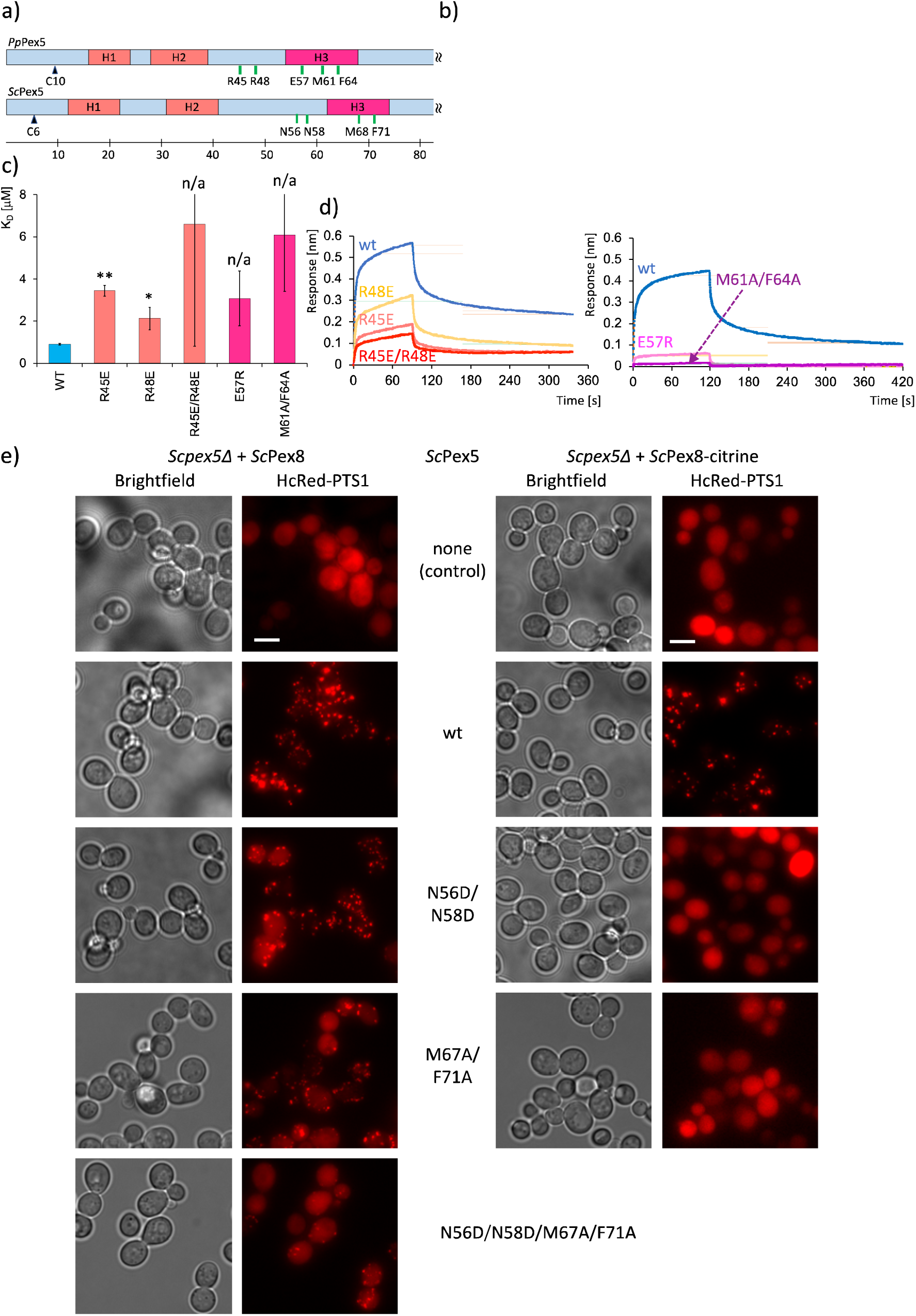
The Pex5 NHB/Pex8 interaction site is essential for cargo translocation. **a**, zoomed scheme of the structural/functional organization of the N-termini of the related *P. pastoris (Pp)* and *S. cerevisiae (Sc)* Pex5 sequences, covering the Pex5 NHB/Pex8 interaction site. Color codes are adopted from previous figures**; b,** Coomassie-stained SDS-PAGE gel of *in vitro* binding assay between Pex5 NHB variants and Pex8. Bands for *Pp*Pex8, GST-*Pp*Pex5 variants, GST (control) and molecular weight markers (M) are labeled; **c,** BLI-Octet quantification of the interactions between GST-*Pp*Pex5 NHB variants and *Pp*Pex8 (see also **Supplementary Figures 16-17**). The data are represented as the mean of N = 3 independent biological replicates ± SD. Statistical significance was determined using a one-tailed, paired Student’s t-test; p=0.0013 < 0.01 (**), p=0.0156 < 0.05 (*), significance not determined (n/a). The dissociation constants K_D_ of GST-*Pp*Pex5 NHB variants R45E/R48E, E57R and M61A/F64A with Pex8 indicated residual binding only, which was below the limits to allow any significance analysis; **d,** representative BLI association/dissociation curves of GST-*Pp*Pex5 NHB variants and Pex8. GST-*Pp*Pex5 NHB variants are shown in related colors to allow their individual identification; **e**, representative microscopy images showing the peroxisomal import of the HcRed-PTS1 fluorescent protein marker in *pex5*Δ cells complemented with different *Sc*Pex5 NHB variants. The cells also expressed either *Sc*Pex8 or *Sc*Pex8-citrine, with the C-terminal PTS1 motif of Pex8 blocked, to demonstrate the dependency of import on the Pex8-Pex5 NTD interaction. Scale bars: 5 µm.

Finally, we mutated several residues in equivalent positions of the predicted H3 helix and the preceding loop of the S. *cerevisiae* AF3 model, including N56, N58, M68 and F71, and assessed their contributions in peroxisomal protein import by epifluorescence microscopy (**Figure 6e**). When S. *cerevisiae pex5*Δ cells were complemented with WT *PEX5*, protein import was restored, as evidenced by the observation of a distinct punctate pattern for peroxisomal matrix protein marker HcRed-PTS1. While expressing the N56D/N58D and M68A/F71A Pex5 double mutants in this background resulted in a partial HcRed-PTS1 import defect, the quadruple mutant (N56D/N58D/M68A/F71A) exhibited a very strong import defect. To further investigate whether the observed import defect was solely attributable to the reduced interaction between the Pex5 NTD and Pex8, we used a *S. cerevisiae pex5*Δ strain expressing a modified Pex8 with a C-terminal Citrine tag. Expression of either N56D/N58D or M68A/F71A Pex5 double mutants completely failed to restore protein import (**Figure 6e**), demonstrating the essential role of the Pex5 NHB interaction with the central HEAT repeat segment of Pex8. In summary, our data performed with purified and proteins in and in yeast cells, consistently demonstrate an essential role of the Pex5 NHB/Pex8 interaction in proper Pex5/Pex8 assembly, as a prerequisite of its essential role to allow import of peroxisomal targets.

## Discussion

Since its initial discovery and characterization more than 25 years ago **[24, 26, 40],** a profound mechanistic understanding of the involvement of the peroxisomal protein Pex8 to the cargo translocation process has been lacking. While the presence of a C-terminal PTS1 Pex5 receptor recognition motif in many Pex8 sequences is suggestive of behaving as a peroxisomal PTS1 cargo, it has become evident that Pex8 – in contrast to most other cargoes – also critically impacts the import of other peroxisomal cargoes regardless of their precise translocation mechanism (**Figure 1) [21, 22, 26].** Earlier studies have indicated that Pex8 has additional functions as a potential cargo releaser and as a connecting unit between the DTM complex and the trimeric peroxisomal RING finger E3-ubiquitin ligase complex, which mono-ubiquitinates Pex5 at an invariant N-terminal cysteine, as the initial step in Pex5 receptor recycling **[18, 19, 21, 23, 41, 42].** Here we revealed that Pex8 functions through an interaction involving a bundle of three helices within the otherwise unfolded Pex5 NTD and the central segment of an irregular HEAT repeat fold of Pex8. This interaction is unique to Pex5 and is independent of other interactions with its folded CTD. Unlike other PTS1 cargoes, the contribution of PTS1-mediated binding of Pex8 into the Pex5 central TPR array groove is only minor.

What could be a possible mechanism behind the functional role of this Pex5 NHB/Pex8 interaction in peroxisomal cargo translocation, and how does it feed into preceding and succeeding steps of this process? The Pex5 NHB interaction with Pex8 is functionally required for proper peroxisomal cargo translocation, in contrast to the functionally dispensable interaction between the Pex5 CTD and the Pex8 PTS1 motif (**Figures 1 and 6).** Therefore, all available evidence points to the Pex5 NHB/Pex8 interaction to be the main driver of Pex5/Pex8 complex formation, effectively avoiding direct competition with other PTS1 cargos for its own translocation into peroxisomes and to become a functional unit to regulate the overall translocation process of other cargoes with diverse receptor recognition motives. The additional Pex5 CTD/Pex8 PTS1 interaction, which increases the binding affinity between Pex5 and Pex8 by up to eightfold (**Figures 2c and 6c),** could play a role in promoting PTS1 cargo release through competition of the Pex8 PTS1 motif with PTS1 cargoes for binding to the Pex5 CTD. However, since most PTS1 cargoes have binding affinities to the Pex5 receptor, that are up to 100-fold higher **[5, 29, 32]**, any direct competition process appears to be unlikely in the absence of the primary Pex5 NHB/Pex8 interaction. Although we did not detect any allosteric interplay between the two Pex5/Pex8 binding sites, sharing these binding sites by the same polypeptide chain enhances the local concentration and could, therefore, shift the underlying thermodynamics of binding to a model favorable for a second Pex5 CTD/Pex8 PTS1 binding site, which could consequently facilitate cargo release.

Furthermore, the interaction between the Pex5 NHB and the central HEAT repeat segment II of Pex8, allowed us to predict a tight assembly with the Pex2/Pex10/Pex12 E3-ubiquitin ligase complex (**Figure 7**). According to the model, the main interaction sites are with HEAT repeats 3, 6, 7, 10 and 12 of Pex8. The interaction with the C-terminal HEAT repeat 12 of Pex8, which includes a the highly conserved WWY motif (residues 700-702) at the outer surface helix of this repeat, involves even two of the three RING finger subunits (Pex2, Pex10) and covers the most widespread interface (**Figure 7a,c, Supplementary Figures 2 and 18**). This model also explains previous data that revealed an essential role of the WWY motif in Pex8-controlled cargo import **[21]**. However, this interface requires the release of PTS1-mediated binding to cargoes, as the binding sites overlap and would lead to steric clashes. This could generate a second event for PTS1 cargo release, even in the absence of bound PTS1-mediated Pex8. Most of the remaining Pex2/Pex10/Pex12 binding sites with Pex8 are distributed over several HEAT repeats of Pex8’s N-terminal and central segments I and II. These sites are close to the Pex5 NHB binding sites on Pex8 but do not overlap. Moreover, the model predicts partial release of the most N-terminal segment of Pex5 into the central tunnel of the Pex2/Pex10/Pex12 trimeric assembly. This would allow for the proper positioning of C10 into the Pex2/Pex10/Pex12 E3-ubiquitin ligase active site, to trigger its mono-ubiquitination. In this model, the first two helices of the NHB, H1 and H2, are part of the released N-terminus of Pex5, whereas helix H3 and the preceding loop connecting to H2 remain bound to Pex8. This is in agreement that all high-confidence interactions in the AF3 models of the Pex5/Pex8 complexes from *P. pastoris* and *S. cerevisiae* are consistently confined to the C-terminal part of the NHB.

**Figure 7.**
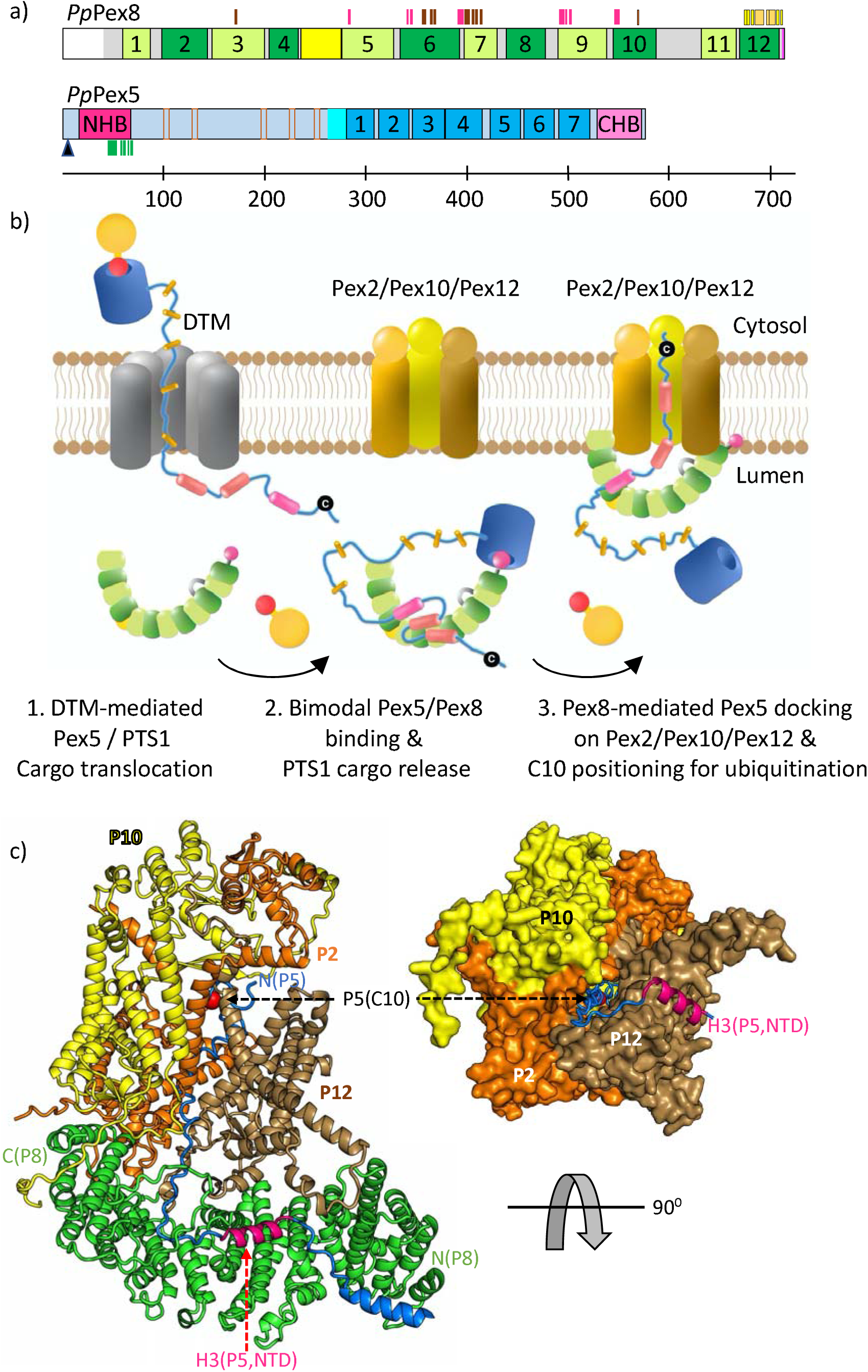
Model of peroxisomal protein import highlighting the role of Pex8 in the Pex5 receptor-mediated cargo translocation cycle. **a**, scheme of the structural/functional organization of the *P. pastoris* Pex5 and Pex8 sequences. Colors are adopted from previous figures. In addition to Pex5/Pex8 interaction sites determined by cryo-EM analysis and integrative modeling (*cf*. Figures 4a **and 5a**), predicted high-confidence sites of the Pex2/Pex10/Pex12 RING finger complex (**Supplementary Table S10**) are indicated by vertical bars in complementary colors of their interaction partners (yellow, orange, brown); **b**, scheme of the key steps of the functional Pex5 receptor cycle, preceding and succeeding bimodal Pex5/Pex8 complex formation (step 2): DTM-mediated Pex5/PTS1 cargo translocation (step 1) and Pex8-mediated Pex5 docking on the Pex2/Pex10/Pex12 RING finger complex, allowing proper positioning of the N-terminal Pex5 cysteine (C10 in *P. pastoris*) into the active site of the RING finger complex to become ubiquitinated (step 3). Whereas the Pex5 NHB H3/Pex8 interaction is required and therefore remains in this step, the Pex5 CTD/Pex8 PTS1 interaction would be incompatible and its disassembly is indicated. Note, that the orientation of the Pex5/cargo complex during DCM-mediated translocation shown is arbitrary. In the absence of its precise knowledge, alternative orientations are possible as well. Pex8 may also interact with the Pex2/Pex10/Pex12 RING finger complex and the peroxisomal membrane in the absence of the Pex5 receptor. In this scheme, the Pex5/Pex8 complex is shown as it was structurally investigated, not directly interacting with the Pex2/Pex10/Pex12 RING finger complex. Protein components and domains are shown in colors adopted from other figures. While the HEAT repeat array of Pex8 is indicated in alternating colors (limon, green), for clarity the Pex5 CTD has been simplified to a hollow cylindrical object in blue. **c,** left panel, cartoon representation of the predicted AF3 model of the pentameric Pex5/Pex8:Pex2/Pex10/Pex12 complex. For the sake of clarity, a truncated model of the trimeric Pex2/Pex10/Pex12 complex is shown, approximately matching a homologous cryo-EM structure of the same complex (PDB entry 7T92). Right panel, combined surface (Pex2/Pex10/Pex12) and cartoon (N-terminal part of the Pex5 NTD only) presentation, illustrating how the N-terminus of Pex5, including NHB helices H1 and H2, disassembles from the Pex8 surface and feeds into a deep groove of the trimeric Pex2/Pex10/Pex12 complex. The predicted location of C10 within the active site of the Pex2/Pex10/Pex12 complex is indicated by a red sphere. Different from previous figures, the N-terminal fragment of Pex5, except NHB H3, is colored in marine, to indicate partial disassembly of the Pex5 NHB from the Pex8 surface when bound to the trimeric Pex2/Pex10/Pex12 complex. For reasons of space, labels P2, P5, P8, P10, and P12 are used as acronyms for Pex2, Pex5, Pex8, Pex10 and Pex12, respectively.

How broadly applicable is this process to different eukaryotic taxa with functional peroxisomes beyond fungal organisms, which universally possess genes coding for Pex5 and Pex8? While the functional characterization of Pex8 from *Arabidopsis thaliana* has been reported recently **[22],** no reliable Pex8 orthologue candidates can be found in metazoan genomes with currently available computational search tools. Whether this is due to a lack of detectability caused by high sequence and structural diversity or if different mechanisms for cargo release and recycling have evolved, remains an open question. Interestingly, support for the latter originates from a recent study, in which a process of proper positioning the N-terminus of Pex5 into the Pex2/Pex10/Pex12 E3-ubiquitin ligase complex appears to be executed by direct interactions with the Pex5 NTD, independent of factors such as Pex8 **[10].** Exploring functional reasons why yeast and plant taxa require such additional factor and why such factor is either unnecessary or is replaced by an unknown factor in metazoan peroxisomal cargo translocation, poses an interesting research question for the future. In summary, our findings have revealed a critical step of Pex5 receptor-driven peroxisomal cargo import, the transition between cargo release as the final step in cargo translocation and receptor recycling (**Figure 7b),** which appears to be more diverse among different taxa than other steps in this cyclic process. This study could help to resolve the remaining puzzle of peroxisomal protein import, in the absence of a high-resolution structure of its translocon.

## Materials and methods

### Cloning and strains

For bacterial protein expression, *P. pastoris* gene fragments were cloned into pETM11 (N-terminal 6xHis tag), pETM14 (N-terminal 6xHis tag), or pETM33 (N-terminal 6xHis-GST tag) vectors. Plasmids harboring Pex8 constructs were expressed in either BL21 LOBSTR *E. coli* strains. Plasmids harboring Pex5 or Pex14 constructs were expressed in Codon RIL *E. coli* strains. For expression in yeast, *S. cerevisiae* gene fragments were cloned into centromeric plasmids Ycplac33 and Ycplac111 **[43],** containing the *LEU2* and *URA3* selectable markers and a *PGK1* terminator. HcRed-PTS1 and Pex5 variants were constitutively expressed from plasmids under the control of *HIS3* and *PEX5* promoters, respectively. Pex5 point mutations were generated by site-directed polymerase chain reaction (PCR) and confirmed by sequencing analysis. The plasmids used in this study are listed in **Supplementary Table 7.**

*S. cerevisiae* strains used in this study are derivatives of BY4741 (*MATA his3*Δ*1 leu2*Δ*0 met15*Δ*0 ura3*Δ*0*) or BY4742 (*MAT*α *his3*Δ*1 leu2*Δ*0 lys2*Δ*0 ura3*Δ*0*) (Euroscarf) and are listed in **Supplementary Table 8**. The pKT239 (pFA6a–link–yECitrine–3HA–SpHIS5) plasmid (Euroscarf) was used as a template for PCR to tag the genomic *PEX8* C-terminally with Citrine-3HA **[44]**. WT and *pex8*Δ strains expressing Pex3 C-terminally tagged with mCherry were crossed with strains containing genomically integrated N-terminal GFP-tagged cargo proteins (*CAT2*, *PCS60*, *FOX2*, *CTA1*, *POX1*, *POT1*) under the control of the *NOP1* promoter **[45].**

### Fluorescence microscopy data acquisition and processing

Yeast strains were grown overnight in synthetic media. For imaging, cultures were diluted 20-fold and incubated at 30 °C for 4 hours. The cells were subsequently transferred to glass-bottom microscope plates (384-well, Matrical Bioscience; 8-well, Ibidi Bioscience), coated with concanavalin A (Sigma-Aldrich). After a 20-minute adherence period, non-adherent cells were removed by washing the wells twice with media. Images shown in Figure 1 were produced at 30 °C using an IXplore SpinSR system (Olympus). This system comprised an Olympus IX83 inverted microscope equipped with a CSU-W1 spinning disk confocal scanner scanning unit (Yokogawa) which included a SoRa module (Yokogawa) that was operated by ScanR software (Olympus). Cells were imaged using an oil immersion objective with 60-fold magnification and a numerical aperture of 1.42. Images were captured with an ORCA-Flash 4.0 camera (Hamamatsu Photonics). Images were acquired in two fluorescence channels.

GFP: excitation at λ = 488 nm and emission at λ = 525/50 nm; Scarlet: excitation at λ = 561 nm and emission at λ = 617/73 nm. For representative images, a single focal plane is shown. The images in Figure 6D were acquired from logarithmically grown yeast cells using a DMi8 inverted live-cell widefield microscope (Leica Microsystems) and analyzed using LAS X Life Science Software (Leica Microsystems). Imaging was conducted using an oil immersion objective with 63-fold magnification and a numerical aperture of 1.40, using a HC PL APO Microscope Objective and Type F immersion oil (Leica Microsystems). Images were captured with a K8 Scientific CMOS camera (Leica Microsystems). Fluorophores were excited by a TANGO LED lamp (Märzhäuser Wetzlar). Images were acquired in a Texas Red fluorescence channel (excitation at λ = 550 nm, emission at λ = 630 nm). Fluorescence images were collected as 0.5 μm *z*-stacks, merged into one plane using Fiji-ImageJ-windows 64-bit (v1.54f) software **[46]** and processed in Photoshop v25.12.1 (Adobe).

### Protein expression and purification

For production of *P. pastoris* Pex5, Pex8 and Pex14 protein constructs (**Supplementary Table 7**), the cells were grown in Terrific-Broth (TB) at 30°C to mid-log phase. Protein expression was induced with 0.5 mM isopropyl-β-D-thiogalactopyranoside (IPTG) overnight at 21°C for Pex8 and at 16°C for Pex5 and Pex14. The cells were harvested at 10,000 *g* for 20 min. The cell pellet was resuspended in a lysis buffer, containing 50 mM HEPES (pH 8.0), 300 mM NaCl, 10 mM imidazole, supplemented with DNAse (1mg/mL=100x stock solution) and EDTA-free protease inhibitor tablets. The cells were lysed by emulsification and the clear lysate was obtained by centrifugation at 4°C and 40,000 g for 40 min. For protein purification, affinity chromatography was performed using a HisTrap HP 5 mL column following the manufacturer’s instructions (Cytiva). Eluted proteins were dialyzed overnight in the lysis buffer without imidazole. To remove affinity tags, Precision 3C or TEV protease was added to the protein during dialysis at a protease/protein ratio of 1:50. The protein samples were again applied to a HisTrap HP column, and the flow-through was collected. For Pex5, an additional ion exchange step was performed using a HiTrap Q HP column (Cytiva). As a final purification step, size exclusion chromatography (SEC) was performed using a HiLoad Superdex 75 or Superdex 200 (16/60) pg column (Cytiva). The Pex5/Pex8 complex for cryo-EM analysis was obtained by mixing purified Pex8 and Pex5 FL at a 1:1 ratio, followed by SEC with a Superdex 200 Increase 10/300 GL column in a buffer containing 50 mM HEPES (pH 7.5), 150 mM NaCl, 0.5 mM Tris(2-carboxyethyl) phosphine (TCEP). The SEC peak fractions were concentrated to ∼1.0 mg/mL and used for subsequent analysis. For selenomethionine (SeMet)-substituted Pex8 production, cells were grown in minimal medium, containing 21.7 g/L medium base MD12-501 (Molecular Dimensions), 5.1 g/L nutrient base MD12-502 (Molecular Dimensions), and 50 mg/L L-SeMet (SERVA). All buffers were supplemented with 1 mM TCEP.

### Biolayer Interferometry analysis

The binding affinities of different Pex5 and Pex8 variants were measured by biolayer interferometry (BLI) using the Octet RED 96 system (FortéBio). Purified 6xHis-GST-tagged Pex5 constructs were loaded onto Penta-His biosensors, pre-equilibrated in assay buffer containing 50 mM HEPES, 150 mM NaCl (pH 7.5) and 0.01% [w/v] Tween-20) for 15 min. Prior to association, a baseline equilibration step was performed for 300 s. Subsequently, the sensors were dipped into wells containing 5 µM Pex8 for 90-120 s (association step), followed by 120-300 s of dissociation time in the same buffer. All experiments were carried out in triplicate at RT. 6xHis-GST was used as a control. The binding data were reference-subtracted and aligned with each other in the Octet Data Analysis software v10.0 (FortéBio), applying a 1:1 binding model. Finally, the binding curves were plotted using either Excel Office 365 (Microsoft) or Prism 10 (GraphPad).

### Isothermal titration calorimetry

All proteins were dialyzed overnight in a buffer containing 50 mM HEPES, 150 mM NaCl (pH 7.5). Appropriate concentrations were obtained by diluting with dialysis buffer. Both buffer and proteins were degassed prior to the ITC experiment. Protein concentrations were at a 1:10 syringe:cell ratio. ITC measurements were conducted at RT using a MicroCal VP-ITC instrument (Malvern Panalytical). Thermograms were integrated by NITPIC **[47].** The data were fitted using SEDPHAT **[48]** and plotted with GUSSI **[47].**

### Structural model prediction

Structural models of the Pex5/Pex8 complexes from *P. pastoris* and *S. cerevisiae* were predicted using AlphaFold 3, using a publicly available web server with default parameters **[49].** Contact predictions were derived and visualized from the distogram, using scripts distributed with gapTrick (DOI: 10.1101/2025.01.31.635911). The same methodology was applied to predict a structural model of the Pex5/Pex8:Pex2/Pex10/Pex12 complex from *P. pastoris*.

### Small Angle X-ray Scattering

All samples were in a solution containing 50 mM HEPES (pH 7.5), 150 mM NaCl, 3% [v/v] glycerol. Small angle X-ray scattering (SAXS) data were measured from a Superdex 200 Increase 10/300 SEC column (Cytiva) eluates at a flowrate of 0.6 mL/min at 23 °C, directly coupled to a 1 mm quartz capillary sample exposure unit under vacuum on beamline P12 at PETRA III (EMBL/DESY, Hamburg, Germany) at 10 keV using a Pilatus-6M photon counting area-detector (Dectris) at a sample-to-detector distance of 3 m **[50],** coupled to a 1260 Infinity II Bio-Inert high-performance liquid chromatography unit (Agilent). The sample injection volume was 90 μL, expect for separate Pex5 FL, for which the volume was 77 μL. Data were recorded as 7200 successive 0.5 s 2D-frames (all samples, except Pex5 FL) or 2400 successive 1.0 s 2D-frames (Pex5 FL), spanning an entire column volume, using the ChemStation software RRID:SCR_015742 (Agilent). Subsequent data reduction by 2D-to-1D azimuthal averaging, considering X-ray absorption by normalizing beam transmission was performed using the *SASFLOW* pipeline **[51].** The final background-corrected SAXS profiles were processed using *CHROMIXS* **[52],** where representative buffer– and sample-scattering data frames were selected, followed by buffer scattering subtraction, absolute scaling relative to the scattering from water, cm^-1^ normalized to beam transmission (except for separate Pex5 FL sample, for which relative scaling in machine units normalized to beam transmission was performed) and averaging. The working s-range was determined using a combination of *AUTORG* (s_min_) [53] and *SHANUM* (s_max_) **[54].** Further details are provided in **Supplementary Table 1**.

Primary SAXS data analysis, including the evaluation of the radius of gyration (*R_g_*), maximum particle dimension (*D_max_*), Porod volume (*V_p_*), and MW estimates for each sample were carried out using modules of the ATSAS 3.0 software package **[55].** The *R_g_* and the extrapolated forward scattering intensity at zero angle *I*(0) were assessed using the Guinier approximation ln(*I*(*s*)) vs. *s*^2^ for *sR_g_* < 1.3 or from the scattering pair distance distribution function *P*(*r*) profiles, using *GNOM* **[56, 57].** The *P*(*r*) profiles also provided estimates of the *D_max_*. The *GNOM* outputs were subsequently used to evaluate the *V_p_* using the ATSAS 3.0 *DATPOROD* tool, in addition to concentration-independent MW estimates using *DATMW* **[58].** Initial models were generated using AF3 and *MultiFoXS* modeling routines **[49, 59]**. For Pex8, residues 1-32 were subsequently removed, to match the purified Pex8 construct. For Pex5, the models were trimmed to the NTD and CTD sequences when experiments were carried out with the respective protein fragments. Model refinement was carried out without applying symmetry constraints and all Pex5/Pex8 complexes were treated in 1:1 stoichiometry. After model refinement (details see below), the fit of each individual model was evaluated with *CRYSOL* using 50 spherical harmonics and constant enabled mode **[60]** to estimate its contribution to the final model ensemble. The quality of the fit was assessed using the reduced χ^2^ test and Correlation-Map (CorMap) *P*-value **[61]**. When *OLIGOMER* was used for refinement, the final model evaluation step was integrated into the refinement procedure. All data and refined models were deposited in the Small Angle Scattering Biological Data Bank (SASBDB**)[62]**, under the accession codes SASDX84 (Pex5 FL) SASDX94 (Pex8), SASDXB4 (Pex5 NTD/Pex8), and SASDXC4 (Pex5 CTD/Pex8), SASDXA4 (Pex5 FL/Pex8).

*SAXS data modeling Pex8.* Residues 233–234 and 587–629 were defined as flexible, to allow for the spatial positioning and optimization to the SAXS data of HEAT-repeat segments I (33–232), II (235–585), insert II (586–629) and HEAT-repeat segment III including the C-terminal PTS1 motif (630–713).

*SAXS data modeling Pex5.* From a volume-fraction weighted limited ensemble of Pex5, the two top-scoring AF3 model predictions were used as inputs into *MultiFoXS* with protein sequence segments defined as flexible. *Model 0*: residues 1–17, 26–30, 39–55, 70–77, 93–128, 138–173, 181–186, 212–219, 237–244, 261–278; *Model 1*: residues 1–17, 26–30, 49–55, 70–77, 94–129, 141–192, 212–219, 239–244, 264–278. *MultiFoXS* was run three times (1x model 0, 2x model 1), and the resulting five-state models were pooled. The fit to the SAXS data of the final volume-fraction weighted limited ensemble was assessed using *OLIGOMER* **[63]**.

*SAXS data modeling Pex8 bound to Pex5 NTD*. The AF3-predicted model of the Pex5 NTD/Pex8 complex was superposed onto the *MultiFoXS*-refined Pex8 model. The interacting Pex5 component from the AF3 prediction (residues 16–91) was subsequently extracted and docked onto the Pex8 *MultiFoXS* model. For subsequent refinement with *CORAL* **[64],** Pex8 was kept as one rigid-body, and four Pex5 sequence segments 16–91, 100–128, 183–214 and 241–263 were defined as separate rigid bodies. The first Pex5 rigid body (residues 16–91) was grouped with the Pex8 model to preserve the AF3-predicted Pex5/Pex8 interface. The remaining Pex5 rigid body segments were connected by flexible dummy-residue linkers, corresponding in length to the number of residues between neighboring Pex5 NTD segments, to sample different spatial positions during 48 times repeated *CORAL* refinement.

*SAXS data modeling Pex8 bound to Pex5 CTD*. The C-terminal PST1 signal sequence (residues 706–713) was removed from the Pex8 template. The resulting Pex8 model was then divided into two rigid bodies, including HEAT repeat segment I (residues 33–233) and the remaining Pex8 sequence (residues 234–705). An AF3-predicted model of the Pex5 CTD bound to the Pex8 PST1 segment was then defined as a third rigid body, and along with the two Pex8 rigid bodies as defined above underwent 3-body *SASREF* rigid-body model refinement to the SAXS data **[65]**. A short-distance constraints of 3.5 Å were imposed between the respective C– and N-termini of each rigid body (Pex8 residues 233/234 and 705/706), in addition to medium-distance constraints of 6.5–12 Å between Pex8 residues 99/276 and 215/283. *SASREF* was run 55 times against the SAXS data to generate a cohort of Pex5 CTD/Pex8 models.

*SAXS data modeling Pex8 bound to Pex5 FL.* The AF3-based *SASREF*-refined Pex5 CTD/Pex8 models (described above) were docked with the predicted Pex5 NTD (residues 16–91)/Pex8 interface. This resulting Pex5 (residues 16–91, CTD)/Pex8 model was further divided into three rigid bodies, which included the N-terminal HEAT repeat segment I of Pex8 (residues 33–233), HEAT repeat segments II and III of the Pex8 sequence (residues 234–705) and the Pex8 PST1 (residues 706–713) bound to the Pex5 CTD. Dummy-residue linkers and short sequence sections for Pex5 residues 100–107, 183–192, 207–214, 241–258 were incorporated into the *CORAL* modeling routines to model the remaining flexible regions of Pex5. A 3.5 Å distance constraint was imposed between the respective C– and N-termini of the Pex8 rigid bodies (residues 233/234 and 705/706), in addition to the of 8–12 Å distance constrains between residues 99/274 and 215/283. *CORAL* was run 123 times to generate a final Pex5/Pex8 model ensemble that fit the SAXS data.

### Crystallization and X-ray structure determination

The purified *P. pastoris* Pex8 was concentrated to a final concentration of 11 mg/mL in a buffer containing 50 mM HEPES (pH 7.5), 150 mM NaCl. Single crystals of Pex8 that diffracted to 2.4 Å resolution were obtained from a solution containing 0.3 M sodium nitrate, 0.3 M disodium hydrogen phosphate, 0.3 M ammonium sulfate, 0.1 M buffer containing 1.0 M HEPES/NaOH, 1.0 M MOPS (pH 7.5) in a 50:50 ratio with a precipitant solution containing 18% precipitant mix of 40% [v/v] ethylene glycol and 20% [w/v] PEG 8,000, and 4% formamide. Data were collected at beamlines P13/P14 at PETRA III (EMBL/DESY, Hamburg, Germany), at 100 K and λ = 0.9762 Å. The diffraction data were indexed, integrated, and scaled using the Aimless suite **[66]**. Experimental phases for structure determination were obtained by single-wavelength anomalous diffraction (SAD), using SeMet-substituted crystals. The diffraction data were analyzed using Phaser_Autosol to obtain experimental phases necessary for building the initial 3D model. This initial model was then used to solve the native structure by molecular replacement (MR) using Phaser-MR software **[67]**. The model was refined iteratively using COOT **[68]** and Refmac5 **[69]** distributed with CCP4 suite version 8.0 **[70].** The final model was validated using MolProbity **[71]** and the PDB deposition server **[72]**.

### Single Particle Cryo-EM analysis

Cryogenic-EM R2/1, Cu 200 mesh perforated carbon grids (Quantifoil) were glow-discharged at 25 mA for 90 s using the GloQube Plus device (Quorum Technologies). Next, 5 µL of the purified Pex8-Pex5 complex at a concentration of 1.0 mg/mL was applied to the glow-discharged grid and blotted for 2.5 s with a blotting force of −5 at 6°C and 100% humidity, followed by draining 2 s and plunge freezing in a 63:37 propane:ethane mixture using a Vitrobot Mark IV device (Thermo Fisher Scientific). Cryo-EM data were acquired on a Titan Krios electron microscope (Thermo Fisher Scientific) operating at 300kV and equipped with a K3 direct electron detector (Gatan). A GIF quantum energy filter (Gatan) with a 20 eV slit width was used for zero-loss filtering. Movies were acquired using EPU software (Thermo Fisher Scientific) in count mode. Each micrograph was fractionated over 60 frames, with a total dose of 64.5 e−/Å2 for 1.8 s. All steps of micrograph image processing were performed using CryoSPARC (v4.4.1) **[73, 74]**. Movies were drift-corrected using patch motion correction. The contrast transfer function (CTF) was performed using patch CTF estimation, and micrographs with a CTF resolution of 5 Å or better were selected for subsequent analysis. Particles were manually picked from a subset of micrographs and were subjected to 2D classification. Selected 2D classes were used for template-based particle picking on the entire dataset. Following multiple rounds of 2D classification, particles with distinct orientations were selected to generate an initial model using *ab initio* reconstruction. This initial model obtained was used as a reference for heterogeneous 3D refinement with three or more classes to assess the level of heterogeneity in the dataset. Selected 3D classes were then subjected to non-uniform 3D refinement. The 3D-refined density map was used for the downstream model-building process.

As the Pex5 FL/Pex8 SAXS input models were composed of rigid domains connected with dummy-atom loop regions, and they were rebuilt to a full-atom representation using gapTrick **[DOI: 10.1101/2025.01.31.635911]** and AF2 **[75].** The resulting models were fitted into cryo-EM reconstructions using tools distributed with the CCPEM suite **[76]**. The models were rigid-body fitted into the cryo-EM reconstruction using MOLREP **[77]** and real-space refined using ISOLDE with self-restraints **[78]**. After this step, flexible loops with no support in map density were removed from the models. The resulting models retained residues 54-66 and 262-576 of Pex5 and residues 33-592 and 629-713 of Pex8. These models were further refined in reciprocal space using Servalcat against unweighted and unsharpened half-maps with jelly-body restraints for 30 cycles **[79]**. All five resulting models have comparable map-model fit score CC_masks from the Phenix suite **[80] (Supplementary Table 9**). A representative model with a CC_mask of 0.75 was further refined using ISOLDE and Servalcat and deposited to the PDB.

### Cross-linking mass-spectrometry analysis

Protein samples were diluted to a final concentration of 21 µM, using crosslinking buffer that contained 50 mM HEPES (pH 7.5) and 150 mM NaCl at 10°C. For each crosslinking reaction 6 µL protein solution was mixed with a second solution of the same protein at the same concentration. After 10 min incubation time, 0.5 µL crosslinking reagent S-SDA (Thermo Fisher Scientific, #26173) dissolved in ultrapure water was added to the protein solution, to achieve a final molar excess of crosslinker over protein of 11.5. N-hydroxy succinimide ester (NHS)-based crosslinking was performed at RT for 30 min. Next, diazirine-based crosslinking was performed using an in-house build LED UV light source (∼ 30 x10^3^ µW/cm^2^) for 20 min on ice. For separating crosslinking reaction products by SDS-PAGE, 3.125 µL of the five-times diluted sample buffer containing 10% [w/v] SDS, 25% [v/v] β-mercaptoethanol, 50% [v/v] glycerol, 1.25 M Tris/HCl (pH 6.8), and 0,01% [w/v] bromophenol blue was added and the mixture was incubated for 8 min at 95°C. Coomassie-stained protein bands were excised and destained by alternating incubations with 10 mM ammonium bicarbonate (AmBic) or 5 mM AmBic / 50% [v/v] ethanol for 10 min at room temperature (RT). Gel slices were dehydrated in 100% ethanol for 10 min at RT, followed by a reduction with 5 mM TCEP in 10 mM AmBic for 45 min at 56°C, and alkylation with 50 mM iodoacetamide (IAA), in 10 mM AmBic for 15 min at RT in the dark. Excess IAA was washed out using 10 mM AmBic and 5 mM AmBic / 50% [v/v] ethanol for 10 min at RT. For subsequent proteolytic digestion, gel slices were dried using a vacuum centrifuge at 30°C for 45 min. Trypsin (Promega) was added at a protease:protein mass ratio of 1:20 and the mixture was digested for 16 h at 37°C. Peptides were eluted using a solution of 50% (v/v) acetonitrile and 50% (v/v) ethanol, and sonicated in an ice-cooled water bath sonicator for 10 min. Eluted peptides were dried in a vacuum centrifuge and either stored at – 80°C or directly resuspended for desalting prior to the LC-MS analysis.

For LC-MS analysis peptides were resuspended in 0.1% (v/v) trifluoroacetic acid and precipitates were pelleted by centrifugation for 6 min at 12,000 rpm at RT). Peptides were separated on a Ultimate^TM^ 3000 RSLCnano system (Thermo Fisher Scientific), equipped with a PepMap^TM^ (C_18_,0.3 mm ID x 5 mm L) precolumn (Thermo Fisher Scientific) and an Acclaim PepMap^TM^ (C_18_, 75 μm ID x 500 mm L, 100 Å pore size) analytical column (Thermo Fisher Scientific), using a binary solvent system consisting of 0.1% [v/v] formic acid (solvent A) and 86% [v/v] acetonitrile/0.1% [v/v] formic acid (solvent B) at 40°C.

Peptides were loaded at 1% solvent B for 3 min at a flow rate of 10 µL x min^-1^ and were eluted by applying a three-step gradient, starting after 4 min with 2% solvent B at a flow rate of 0.3 µL x min^-1^ over 24% solvent B at 67 min, 31% at 80 min, and reaching 47% solvent B after 91 min. Subsequently, the column was flushed with 95% solvent B for 5 min.

Finally, the column was equilibrated for 20 min at 1% solvent B with a flow rate of 0.35 µL x min^-1^. Separated peptides were measured on a Q Exactive Plus mass spectrometer (Thermo Fisher Scientific), equipped with a nano electrospray ion source and a stainless steel emitter, using a data-dependent acquisition mode with a scan range of *m/z* 400 to 1,450 at a maximum resolution of 140,000 at *m/z* 400, an automatic gain control (AGC) of 3 x 10^6^ ions and a maximum fill time of 50 ms for MS^1^ scans. The 10 most intense precursor ions with charge states between 2+ and 8+ were selected for higher-energy collisional dissociation (HCD) with stepped normalized collision energies of 24%, 28% and 30%. MS^2^ data were acquired within a scan range of *m/z* 200 to 2,000 at a resolution of 70,000, with an AGC of 5 x 10^4^ and a maximum fill time of 120 ms. A dynamic exclusion time of 30 s was set for already fragmented precursor ions.

Crosslinked peptides were identified using the software pLink2 (version 2.3.9) **[81]**. Searches were performed against the forward and reverse sequences of protein samples, including tryptic peptides 5-60 amino acids in length and allowing a maximum of three missed cleavages. Oxidation of methionine was set as variable and carbamidomethylating of cysteine was treated as a fixed modification. Lysine, tyrosine, serine, threonine and the N-terminus were set as one crosslinked site, and any other residue as second crosslinked site. The mass tolerance of the search of MS^1^ and MS^2^ spectra was set to 20 ppm, and identified MS^1^ spectra were subsequently filtered by a mass tolerance of 10 ppm and a false discovery rate (FDR) of 5% at peptide-spectrum match (PSM) level. pLink2 results were subsequently filtered by the following criteria: PSMs having E-values above 1 x 10^-2^ or comprising adjacent tryptic peptides of the same protein were rejected. Additionally, crosslinks covering the protein N-terminus were omitted. Only crosslinked residue pairs that were identified in 3 out 4 replicates were retained for further interpretation. The resulting crosslinking data were then visualized using XiNET (version 2.0.0) **[82].**

### Integrative structural model building and refinement

To generate complete structural models of the Pex5/Pex8 complex, we incorporated available distance restraints derived from XL-MS analysis into the gapTrick-rebuilt SAXS models (for further details, see above). C_α_/C_α_ atom distance restrains of 25 Å were generated for all residue pairs, for which crosslinks were identified in XL-MS analysis, in ISOLDE **[78].** The resulting restrained models were further refined by real-space molecular dynamics in ISOLDE, leading to a final model ensemble, in which all XL-MS-derived distances were within a permissive range of <25 Å.

### GST-Pull down assays

For *in vitro* binding assays, a small-scale expression and purification of GST-tagged Pex5 NTD variants were performed using Glutathione-Sepharose 4B beads (Cytiva). Cells expressing GST-Pex5 NTD variants were lysed in a buffer containing 50 mM HEPES (pH 7.5), 150 mM NaCl, and 0.01% [w/v] Tween-20. The lysate was then incubated with Glutathione-Sepharose 4B beads for 1 h at 4°C through shaking at 1,000 rpm. The beads were washed to remove unbound proteins, and a total of 50 µL of the respective binding partner at a concentration of 5 µM was added to the GST-Pex5 NTD-bound beads, followed by incubation for 45 min at 4°C through shaking at 1,000 rpm. After washing to remove unbound proteins, the bound proteins were eluted by adding fourfold diluted 1 x NuPAGE^TM^ lithium dodecyl sulfate (LDS) Sample Buffer (NP0008, Thermo Fisher Scientific) and boiling for 5 min at 75°C.

### Statistics and reproducibility

The legends of the respective figure panels provide relevant information on statistical data analysis. For plotting graphs and statistical analysis, GraphPad Prism 10.3.1 software was used. Statistical analysis was performed using a paired t-test. ****p<0.0001; ***P<0.005; **p<0.01; *p<0.05; non-significant (ns) p>0.05.

## Acknowledgements

We thank for access to and technical support at the EMBL Hamburg beamlines P12, P13 and P14, operated at the PETRA III storage ring (DESY, Hamburg, Germany), the EMBL Hamburg Sample Preparation and Characterization (SPC) Facility, and the cryo-EM and Advanced Light and Fluorescence Microscopy facilities at the Centre for Structural Systems Biology (CSSB), Hamburg. This work was supported by the Deutsche Forschungsgemeinschaft (DFG, German Research Foundation) within the research unit FOR1905 “PerTrans” to M.W. (WI 1058/9-2) and to BW (Project ID 219314758) and the Research Training Group (RTG) 2202 (Project-ID 278002225) to BW. Work in the MS and EZ labs is supported by an Israeli Science Foundation (ISF) grant 914/22. We thank Krisztian Fodor, Daniel Passon, Indra Bekere, Nabil Hanna, Morlin Milewski, Jérôme Bu rgi, and Amal Hassan for their contributions during the early stages of this project at the EMBL Hamburg Unit, Hamburg, Germany. We also thank Manjeet Kumar and Toby Gibson from the EMBL Molecular Systems Biology Unit (formerly the Structural and Computational Biology Research Unit), Heidelberg, Germany, for their computational work during the early phase of this project. Finally, we thank Dr. Ewald Hettema, University of Sheffield, UK, for providing yeast vectors Ycplac33 and Ycplac111.

## Author contributions

L.E., D.W., G.C., B.W., E.Z. and M.W. designed the experiments. L.E., D.W., Y.D. and E.M. performed the experiments. L.E., D.W., G.C., C.M.J., E.M., E.Z. and M.W. analyzed the data. L.E. and M.W. wrote the paper, including contributions on specific sections by D.W., G.C., C.M.J. and E.Z. M.S., B.W. and M.W. supported the project.

