## Supplement for "Structure of Pex8 in complex with peroxisomal receptor Pex5 reveals its essential role in peroxisomal cargo translocation"

**SUPPLMENTARY TABLES**

### Supplementary Table 1a. SAXS data collection and processing

| **Samples** | | **Pex5 FL** | **Pex8** |
| --- | --- | --- | --- |
| Deposition (SASDBD) | | SASDX84 | SASDX94 |
| **Size Exclusion Chromatography SEC-SAXS and SAS data collection** | | | |
| Sample concentration (mg/mL) | 8.0 | | 4.0 |
| Measured *s*-range (*s_min_*–*s_max_* nm^-1^) | 0.030–7.26 | | 0.024–7.39 |
| Working *s*-range (*s_min_*–*s_max_* nm^-1^) | 0.052–3.83 | | 0.057–4.06 |
| Number of frames used for averaging | 59 | | 46 |
| **SAS-derived structural parameters** | | | |
| *Guinier Analysis R*_g_ *correlation/stability through SEC-elution peak*. | | | |
| Mean *R*_g_ ± σ (nm) | 4.85 (0.11) | | 3.65 (0.07) |
| Median *R*_g_ (nm) | 4.85 | | 3.63 |
| *R*_g_ min-max range (nm) | 4.61–5.28 | | 3.53−3.81 |
| *Guinier Analysis (final scaled and averaged data)* | | | |
| *I*(0) ± σ (cm^-1^) | 2351.1 ± 4.2 | | 0.01741 ± 0.0001 |
| *R*_g_ ± σ (nm) | 5.03 ± 0.02 | | 3.61 ± 0.02 |
| *min < sR_g_* < *max* limit (nm)^a^ | 0.29–1.11 (11–70) | | 0.21–1.30 (13–122) |
| Pearson correlation coefficient R^2^ | 0.9964 | | 0.9923 |
| *P(*r*) analysis* | Pex5 | | Pex8 |
| *I*(0) ± σ (cm^-1^) | 2377 ± 6 | | 0.01762 ± 0.0001 |
| *R*_g_ ± σ (nm) | 5.37 ± 0.03 | | 3.76 ± 0.01 |
| *D*_max_ (nm) | 21.4 | | 13 |
| *s*-range for fit (nm^-1^) | 0.052–3.35 | | 0.057–3.36 |
| *P*(*r*) fit assessment *χ*^2^ (CorMap-*P*) | 0.973 (0.44) | | 0.993 (0.134) |
| **Scattering particle size** | | | |
| Porod volume, *V_p_* (nm^3^) | 131 | | 117 |
| MW calculated from sequence (kDa) | 65.2 | | 78.0 |
| MW range from SAXS/*P*(*r*) > 0.9 confidence) (kDa) | 80 (73–87) | | 74 (67–75) |
| **Modelling** | | | |
| *s*-range for fit (nm^-1^) | 0.052–3.83 | | 0.057–4.06 |
| Number of model reconstructions | 9-member limited ensemble | | 1 |
| *χ*^2^ (CorMap *P*-values for fit) | 1.02 (0.486)^c^ | | 0.99 (0.136)^d^ |

^a^Data point range is relative to the full data-point/*s*-range of the measured SAXS profile.

### Supplementary Table 1b. SAXS data collection and processing summary

| **Samples** | **Pex5 FL/Pex8** | | **Pex5 NTD/Pex8** | **Pex5 CTD/Pex8** | |
| --- | --- | --- | --- | --- | --- |
| Deposition (SASDBD) | SASDXA4 | | SASDXB4 | SASDXC4 | |
| **Size Exclusion Chromatography SEC-SAXS and SAS data collection** | | | | | |
| Sample concentration (mg/mL) | 7.5 | 5.2 | | | 5.8 |
| Measured *s*-range (*s_min_*–*s_max_* nm^-1^) | 0.024–7.40 | 0.024–7.40 | | | 0.024–7.40 |
| Working *s*-range (*s_min_*–*s_max_* nm^-1^) | 0.060–4.06 | 0.082–4.00 | | | 0.071–3.50 |
| Number of frames used for averaging | 86 | 87 | | | 63 |
| **SAS-derived structural parameters** | | | | | |
| *Guinier Analysis R*_g_ *correlation/stability through SEC-elution peak*. | | | | | |
| Mean *R*_g_ ± σ (nm) | 5.14 ± 0.05 | 4.53 ± 0.11 | | | 4.20 ± 0.04 |
| Median *R*_g_ (nm) | 5.15 | 4.57 | | | 4.20 |
| *R*_g_ min-max range (nm) | 5.01–5.25 | 4.25−4.73 | | | 4.09–4.28 |
| *Guinier Analysis (final scaled and averaged data)* | | | | | |
| *I*(0) ± σ (cm^-1^) | 0.05327 ± 0.0001 | 0.02796 ± 0.0001 | | | 0.03575 ± 0.0001 |
| *R*_g_ ± σ (nm) | 5.15 ± 0.01 | 4.56 ± 0.02 | | | 4.21 ± 0.01 |
| *min < sR_g_* < *max* limit (nm)^a^ | 0.31–1.27 (14–81) | 0.37–1.29 (22–94) | | | 0.30–1.30 (18–103) |
| Pearson correlation coefficient R^2^ | 0.9985 | 0.9961 | | | 0.9974 |
| *I*(0) ± σ (cm^-1^) | 0.05407 ± 0.0001 | 0.02832 ± 0.0001 | | | 0.03591 ± 0.0001 |
| *R*_g_ ± σ (nm) | 5.46 ± 0.03 | 4.80 ± 0.02 | | | 4.31 ± 0.01 |
| *D*_max_ (nm) | 23.5 | 19 | | | 14.5 |
| *s*-range for fit (nm^-1^) | 0.060–3.50 | 0.082–3.50 | | | 0.071–3.50 |
| *P*(*r*) fit assessment *χ*^2^ (CorMap-*P*) | 0.961 (0.993) | 1.13 (0.072) | | | 1.05 (0.912) |
| **Scattering particle size** |  |  | | |  |
| Porod volume, *V_p_* (nm^3^) | 247 | 177 | | | 141 |
| MW calculated from sequence (kDa) | 143.2 | 109.9 | | | 114.1 |
| MW range from SAXS/*P*(*r*) > 0.9 confidence) (kDa) | 157 (142–163) | 119 (111–127) | | | 101 (92–107) |
| **Modelling** | | | | | |
| *s*-range for fit (nm^-1^) | 0.060–4.06 | 0.082–4.00 | | | 0.071–3.50 |
| Number of model reconstructions | 6 | 5 | | | 5 |
| *χ*^2^ (CorMap *P*-values for fit) | 0.97–1.03 (0.907–0.07) | 1.09–1.11 (0.689–0.251) | | | 1.09–1.13 (0.906–0.135) |

^a^ Data point range is relative to the full data-point/*s*-range of the measured SAXS profile .

### Supplementary Table 2a. High probability contacts extracted from AF3 structure prediction of *P.* *pastoris* Pex5/Pex8 complex.

| ***Pp*Pex5** | ***Pp*Pex8** |  |  |  |  |  |  |
| --- | --- | --- | --- | --- | --- | --- | --- |
| R45 | I549 |  |  |  |  |  |  |
| M46 | Y498 | I549 |  |  |  |  |  |
| M47 | E494 | H497 | Y498 | L501 | Q546 | I549 | A550 |
| R48 | S491 | E494 | S495 | Y498 |  |  |  |
| N49 | S491 |  |  |  |  |  |  |
| E50 | G392 | T393 | G394 |  |  |  |  |
| S51 | G392 | T393 | G394 |  |  |  |  |
| T52 | G392 | G394 | G395 |  |  |  |  |
| M53 | I391 | G392 | T393 | G394 | G395 | F396 |  |
| E57 | I391 | G392 |  |  |  |  |  |
| Q60 | R340 |  |  |  |  |  |  |
| M61 | Y344 | F396 |  |  |  |  |  |
| F64 | N282 | Q341 | Y344 |  |  |  |  |
| M65 | N282 | Y344 |  |  |  |  |  |
| Q67 | N282 |  |  |  |  |  |  |
| E326 | K712 |  |  |  |  |  |  |
| E328 | K712 |  |  |  |  |  |  |
| I354 | L713 |  |  |  |  |  |  |
| I357 | L713 |  |  |  |  |  |  |
| N358 | K712 | L713 |  |  |  |  |  |
| V430 | L713 |  |  |  |  |  |  |
| Y433 | L713 |  |  |  |  |  |  |
| N460 | L713 |  |  |  |  |  |  |
| R461 | L713 |  |  |  |  |  |  |
| G463 | A711 |  |  |  |  |  |  |
| A464 | A711 | K712 | L713 |  |  |  |  |
| A467 | A711 |  |  |  |  |  |  |
| N468 | T710 | A711 |  |  |  |  |  |
| A475 | A711 |  |  |  |  |  |  |
| R491 | K712 |  |  |  |  |  |  |
| Y494 | K712 |  |  |  |  |  |  |
| N495 | A711 | K712 |  |  |  |  |  |
| V498 | S709 | T710 | A711 |  |  |  |  |
| S499 | A711 |  |  |  |  |  |  |
| T544 | S709 |  |  |  |  |  |  |

### Supplementary Table 2b. High probability contacts extracted from AF3 structure prediction of *S. cerevisiae* Pex5/Pex8 complex.

| ***Sc*Pex5** | ***Sc*Pex8** |  |  |  |  |  |  |
| --- | --- | --- | --- | --- | --- | --- | --- |
| I49 | T432 | I435 | T436 | L479 | H483 |  |  |
| S50 | N385 | T432 | N385 | T432 |  |  |  |
| A52 | L479 |  |  |  |  |  |  |
| F53 | E428 | H431 | T432 | I435 | Q476 | L479 |  |
| I54 | E425 | E428 | T432 |  |  |  |  |
| S55 | I424 | E428 | S473 |  |  |  |  |
| N56 | E425 |  |  |  |  |  |  |
| V57 | V333 |  |  |  |  |  |  |
| N58 | T327 | Q328 | G331 | T332 | V333 |  |  |
| A59 | G331 | V333 |  |  |  |  |  |
| I60 | I330 | G331 | T332 | V333 | G334 | F335 |  |
| S61 | G331 |  |  |  |  |  |  |
| N64 | Q284 | I330 | G331 |  |  |  |  |
| N67 | F281 | Q284 | N285 |  |  |  |  |
| M68 | I288 | F335 |  |  |  |  |  |
| F71 | L234 | N235 | A238 | N285 | I288 |  |  |
| I72 | Q185 | N235 |  |  |  |  |  |
| G74 | N184 | Q185 | N235 | H236 |  |  |  |
| E75 | N235 |  |  |  |  |  |  |
| P76 | T232 |  |  |  |  |  |  |
| L77 | S231 | T232 | N233 | L234 | N235 | F281 | N285 |
| I78 | F281 |  |  |  |  |  |  |
| D79 | S231 | T232 | T232 |  |  |  |  |
| E361 | K588 |  |  |  |  |  |  |
| E363 | K588 |  |  |  |  |  |  |
| I389 | L589 |  |  |  |  |  |  |
| I392 | L589 |  |  |  |  |  |  |
| N393 | K588 | L589 |  |  |  |  |  |
| L465 | L589 |  |  |  |  |  |  |
| Y468 | L589 |  |  |  |  |  |  |
| N495 | L589 |  |  |  |  |  |  |
| G496 | L589 |  |  |  |  |  |  |
| G498 | S587 |  |  |  |  |  |  |
| A499 | S587 | K588 | L589 |  |  |  |  |
| S500 | L589 |  |  |  |  |  |  |
| A502 | S587 |  |  |  |  |  |  |
| N503 | S586 | S587 |  |  |  |  |  |
| A510 | S587 |  |  |  |  |  |  |
| R526 | K588 |  |  |  |  |  |  |
| Y529 | K588 |  |  |  |  |  |  |
| N530 | S587 | K588 |  |  |  |  |  |
| S534 | S587 |  |  |  |  |  |  |

### Supplementary Table 3. Cryo-EM data collection and processing summary

| **Accession code** | **EMD** |
| --- | --- |
| **Data collection and processing** | |
| Microscope | TFS Krios |
| Detector | Gatan K3 counting mode |
| Software | EPU Thermo Fisher Scientific |
| Acceleration voltage (kV) | 300 |
| Pixel size (Å/pixel) | 0.68 |
| Energy filter slit width | 20 eV |
| Magnification | 13,0000 |
| Defocus range (μm) | −0.75 to −2.5 |
| Number of frames | 60 |
| Exposure time (s) | 1.8 |
| Total exposure (e-/Å^2^) | 64.5 |
| Total micrographs collected | 31235 |
| Total micrographs used | 28198 |
| Data processing software | CryoSPARC v4.4.1 |
| Particles after 2D classification | 476052 |
| Particles in final 3D reconstruction | 85869 |
| Symmetry imposed | C1 |
| Map sharpening B-factor | 0 |
| Masked resolution at 0.5/0.143 FSC (Å) | 4.5 |
| Local resolution range (Å) | 4.1 – 9.3 |
| **Structure refinement** | |
| Model to map fit (FSC average) | 0.86 |
| Ramachandran favored (%) | 98 |
| Ramachandran allowed (%) | 2 |
| Ramachandran outliers (%) | 0 |
| RAMA-Z score | -0.53 |
| Poor rotamers (%) | 0.7 |
| Clash score | 3 |
| MolProbity score | 1.09 |
| CC_mask | 0.66 |

### Supplementary Table 4. X-ray data structure determination for Pex8.

| **Accession code (PDB)** | 8RQT |
| --- | --- |
| **Data Collection and processing** |  |
| Wavelength (nm) | 0.097 |
| Molecules / asymmetric unit | 1 |
| Space group | P 2_1_ 2_1_ 2_1_ |
| Unit cell dimensions (Å) | 81.2, 87.8, 153.1 |
| Solvent content (%) | 63.5 |
| Resolution range (Å) | 47.03 - 2.41 (2.50-2.41) |
| Unique reflections | 43103 (4486) |
| Completeness (%) | 100.00 (100.00) |
| I/σ(I) | 10.00 (0.70) |
| Multiplicity | 13.20 (13.80) |
| R_merge_ (%) | 19.5 (50.5) |
| CC_1/2_ | 1.00 (0.41) |
| **Structure refinement** | |
| R_work,_ R_free_ | 0.213, 0.257 |
| Number non-hydrogen atoms | 5082 |
| Protein | 5016 |
| Solvent | 66 |
| r.m.s.d. (bonds) | 0.0 |
| r.m.s.d. (angles) | 0.1 |
| Ramachandran favored (%) | 98.0 |
| Ramachandran allowed (%) | 2.0 |
| Ramachandran outliers (%) | 0.0 |
| Rotamer outliers (%) | 2.0 |
| Clash score | 2.0 |
| Average B-factor | 77 |
| Number of TLS groups | 1 |

### Statistics for the highest resolution shell are in parentheses.

### Supplementary Table 5. HEAT repeat geometry

|  | **Pex8 (crystal structure)** | | **Pex8 (EM structure)** | |
| --- | --- | --- | --- | --- |
| **Angles between helical axes from neighboring HEAT repeats (n/n+1)** | **Inner helices (deg)** | **Outer Helices (deg)** | **Inner helices (deg)** | **Outer Helices (deg)** |
| 1/2 | 13.4 | 35.5 | 12.3 | 36.2 |
| 2/3 | 33.3 | 24.9 | 37.4 | 20.8 |
| 3/4 | 25.4 | 11.9 | 29.5 | 16.1 |
| 4/5 | 91.4 | 95.5 | 86.2 | 83.1 |
| 5/6 | 10.0 | 16.8 | 9.4 | 13.4 |
| 6/7 | 50.0 | 11.4 | 48.0 | 12.0 |
| 7/8 | 49.6 | 41.4 | 49.8 | 33.4 |
| 8/9 | 16.4 | 12.9 | 15.6 | 16.4 |
| 9/10 | 16.8 | 4.6 | 14.7 | 7.9 |
| 10/11 | 30.2 | 4.0 | 32.4 | 7.7 |
| 11/12 | 16.8 | 31.3 | 9.3 | 30.0 |
| **Summary** | **353.3** | **290.2** | **344.6** | **277** |

**Supplementary Table 6a: Pex5 FL/Pex8 crosslinkers**

| **No.** | ***Pp*Pex5 FL** | **Residue** | ***Pp*Pex8** | **Residue** |
| --- | --- | --- | --- | --- |
| 1 | NGEFQQGNQR | 36 | LWDCVVGTNKYDPQK | 688 |
| 2 | AGIYGGGR | 158 | LKNEQNVALLTDFFLR | 71 |
| 3 | AGIYGGGR | 158 | SIKGSDTAVAQLLQR | 178 |
| 4* | AGIYGGGR- | 161 | FKVSQPTVPFYR | 117 |
| 5 | TQEPKTK | 215 | GRDDVELYPDHSR | 170 |

(*) Pex5 FL / Pex8 only

**Supplementary Table 6b: Pex5 FL/Pex8 integrative model crosslinker distances [Å]**

|  | **Pex5** | **Pex8** | **yPx8517** | **rPx8512** | **rPx8510** | **oPx8512** | **qPx8515** | **Average** |
| --- | --- | --- | --- | --- | --- | --- | --- | --- |
| 1 | N36 | K688 | 23.2 | 22.5 | 22.6 | 22.0 | 22.4 | 22.5 |
| 2 | A158 | K71 | 20.2 | 19.0 | 20.1 | 24.2 | 21.6 | 21.0 |
| 3 | A158 | K178 | 24.4 | 23.6 | 21.8 | 19.9 | 22.2 | 22.4 |
| 4 | Y161 | K117 | 22.1 | 18.9 | 15.2 | 21.1 | 16.3 | 18.7 |
| 5 | K215 | Y170 | 22.5 | 20.5 | 20.6 | 23.4 | 23.7 | 22.1 |

### Supplementary Table S7. The plasmids used in this study.

| **Vector backbone** | **Insert (residue range)** | **Acronym** | **Tag** |
| --- | --- | --- | --- |
| pETM11 | *Pp*Pex5 (1-576) | FL | N’ 6xHis-TEV |
| pETM33 | - | Control | N’ 6xHis-GST_3C |
| pETM33 | *Pp*Pex5 (1-276) | NTD | N’ 6xHis-GST_3C |
| pETM33 | *Pp*Pex5 (1-276) R35E | NTD | N’ 6xHis-GST_3C |
| pETM33 | *Pp*Pex5 (1-276) R45E | NTD | N’ 6xHis-GST_3C |
| pETM33 | *Pp*Pex5 (1-276) R48E | NTD | N’ 6xHis-GST_3C |
| pETM33 | *Pp*Pex5 (1-276) R58E | NTD | N’ 6xHis-GST_3C |
| pETM33 | *Pp*Pex5 (1-276) R45/48E | NTD | N’ 6xHis-GST_3C |
| pETM33 | *Pp*Pex5 (1-276) E57R | NTD | N’ 6xHis-GST_3C |
| pETM33 | *Pp*Pex5 (1-276) M61A/F64A | NTD | N’ 6xHis-GST_3C |
| pETM33 | *Pp*Pex5 (1-276) F64W | NTD | N’ 6xHis-GST_3C |
| pETM33 | *Pp*Pex5 (1-276) M61W/F64W | NTD | N’ 6xHis-GST_3C |
| pETM14 | *Pp*Pex5 (259-576) | CTD | N’ 6xHis-3C |
| pETM14 | *Pp*Pex8 (33-713) |  | N’ 6xHis-3C |
| pETM14 | *Pp*Pex8 ∆AKL (33-710) | ∆PTS1 | N’ 6xHis-3C |
| pETM14 | *Pp*Pex14 (1-124) | NTD | N’ 6xHis-3C |
| Ycplac33 | *Hc*Red-PTS1 |  |  |
| Ycplac111 | Control plasmid |  |  |
| Ycplac111 | *Sc*Pex5 (1-612) |  | N’ 3xHA |
| Ycplac111 | *Sc*Pex5 (1-612) K46D |  | N’ 3xHA |
| Ycplac111 | *Sc*Pex5 (1-612) N56D |  | N’ 3xHA |
| Ycplac111 | *Sc*Pex5 (1-612) N58D |  | N’ 3xHA |
| Ycplac111 | *Sc*Pex5 (1-612) N56/58D |  | N’ 3xHA |
| Ycplac111 | *Sc*Pex5 (1-612) M68A/F71A |  | N’ 3xHA |
| Ycplac111 | *Sc*Pex5 (1-612) N56D/N58D/M68A/F71A |  | N’ 3xHA |
| pKT239 | yeCitrine |  | C’ 3xHA |

### Supplementary Table 8. Yeast strains used in this study.

| **Strain and genotype** | **Source** |
| --- | --- |
| BY4741 MATa *his3Δ1 leu2Δ0 met15Δ0 ura3Δ0* | Euroscarf |
| BY4742 MATα *his3Δ1 leu2Δ0 lys2Δ0 ura3Δ0* | Euroscarf |
| BY4742 *pex5Δ::kanMX4* | Euroscarf |
| BY4741 *pex5Δ::kanMX4 PEX8::PEX8-citrine-his3MX6* | This study |
| yMSxxxx *PEX3::PEX3-mCherry-natMX6* |  |
| yMSxxxx *NOP1pr*-*GFP-CAT2* | This study |
| yMSxxxx *NOP1pr*-*GFP-PCS60* | This study |
| yMSxxxx *NOP1pr*-*GFP-FOX2* | This study |
| yMSxxxx *NOP1pr*-*GFP-CTA1* | This study |
| yMSxxxx *NOP1pr*-*GFP-POX1* | This study |
| yMSxxxx *NOP1pr*-*GFP-POT1* | This study |
| yMSxxxx *pex8Δ::kanMX4 NOP1pr*-*GFP-CAT2* | This study |
| yMSxxxx *pex8Δ::kanMX4 NOP1pr*-*GFP-PCS60* | This study |
| yMSxxxx *pex8Δ::kanMX4 NOP1pr*-*GFP-FOX2* | This study |
| yMSxxxx *pex8Δ::kanMX4 NOP1pr*-*GFP-CTA1* | This study |
| yMSxxxx *pex8Δ::kanMX4 NOP1pr*-*GFP-POX1* | This study |
| yMSxxxx *pex8Δ::kanMX4 NOP1pr*-*GFP-POT1* | This study |
| BY4742 *pex5Δ::kanMX4* | Euroscarf |
| BY4742 *pex5Δ::kanMX4 PEX8::PEX8-citrine-his3MX6* | This study |

### Supplementary Table 9. Integrative model refinement statistics.

| **Model ID** | **SAXS** | **Cryo-EM** | **XL-MS** |
| --- | --- | --- | --- |
| **Validation** | **X**2** | **CC_mask** | **XLs < 25 Å** |
| 141 [yPx8517] | 1.03 | 0.72 | 5 |
| 131 [rPx8512] | 0.98 | 0.74 | 5 |
| 143 [rPx8510]* | 0.97 | 0.75 | 5 |
| 142 [oPx8512] | 1.00 | 0.74 | 5 |
| 149 [qPx8515] | 0.99 | 0.73 | 5 |

### Integrative modeling of Pex5/Pex8 complexes. Pex5/Pex8 structural models predicted by AF3 (*cf.* Supplementary Figure 3) were used to create an ensemble of 5 Pex5/Pex8 models fitting the SAXS data of Pex5 FL in complex with Pex8 (*cf*. Supplementary Figure 8e). These models served as input for independent refinement against the cryo-EM density map (Figure 4b-c, Supplementary Figure 9) and distance restraints derived from crosslinking mass spectrometry (XL-MS) analysis (Figure 5a-b, Supplementary Figure 12). The resulting model ensembles are shown in Supplementary Figures 13-14.

### Table S10. High probability contacts extracted from AlphaFold3 structure prediction of *P.* *pastoris* Pex5-Pex8-E3 ubiquitin ligase (Pex2/Pex10/Pex12) complex

| ***Pp*Pex5** | ***Pp*Pex8** |  |  |  |  |  |  |
| --- | --- | --- | --- | --- | --- | --- | --- |
| R45 | I549 |  |  |  |  |  |  |
| M46 | Y498 | I549 |  |  |  |  |  |
| M47 | E494 | H497 | Y498 | L501 | Q546 | I549 | A550 |
| R48 | S491 | E494 | S495 | Y498 |  |  |  |
| N49 | S491 | E494 |  |  |  |  |  |
| E50 | G392 | T393 | G394 |  |  |  |  |
| S51 | G392 | T393 | G394 |  |  |  |  |
| T52 | G392 | G394 | G395 | G395 |  |  |  |
| M53 | I391 | G392 | G394 | G395 | F396 |  |  |
| E57 | I391 | G392 |  |  |  |  |  |
| Q60 | R340 |  |  |  |  |  |  |
| M61 | Y344 | F396 |  |  |  |  |  |
| F64 | N282 | Q341 | Y344 |  |  |  |  |
| M65 | Y344 | P398 |  |  |  |  |  |
| Q67 | N282 |  |  |  |  |  |  |
| ***Pp*Pex5** | ***Pp*Pex2/ *Pp*Pex10/*Pp*Pex12** | | | | | | |
| N15 | Y38 (P12) |  |  |  |  |  |  |
| P16 | L31 (P12) | P34 (P12) | S35 (P12) | Y38 (P12) |  |  |  |
| L17 | S35 (P12) | Y38 (P12) | 39 (P12) | 282 (P12) |  |  |  |
| 1L7 | Y38 (P12) | I39 (P12) | 282 (P12) |  |  |  |  |
| Q19 | L31 (P12) |  |  |  |  |  |  |
| F20 | L28 (P12) | L31 (P12) | S35 (P12) |  |  |  |  |
| T21 | K283 (P12) | E286 (P12) |  |  |  |  |  |
| H23 | E27 (P12) | L28 (P12) | L31 (P12) |  |  |  |  |
| T24 | I279 (P12) |  |  |  |  |  |  |
| D27 | F221 (P10) |  |  |  |  |  |  |
| T28 | S24 (P12) | F221 (P10) |  |  |  |  |  |
| S29 | S218 (P10) | F221 (P10) |  |  |  |  |  |
| L30 | D19 (P10) | L23 (P10) |  |  |  |  |  |
| Q31 | F20 (P12) |  |  |  |  |  |  |
| ***Pp*Pex8** | ***Pp*Pex2/ *Pp*Pex10/*Pp*Pex12** | | | | | | |
| K117 | I243 (P12) |  |  |  |  |  |  |
| V167 | I243 (P12) |  |  |  |  |  |  |
| E168 | R244 (P12) |  |  |  |  |  |  |
| Y170 | T249 (P12) | E250 (P12) | Y253 (P12) |  |  |  |  |
| D172 | T247 (P12) |  |  |  |  |  |  |
| T355 | R113 (P12) |  |  |  |  |  |  |
| F357 | T96 (P12) | I97 (P12) |  |  |  |  |  |
| F358 | H94 (P12) | I97 (P12) | R113 (P12) | R114 (P12) |  |  |  |
| D361 | H94 (P12) | T96 (P12) | K206 (P12) |  |  |  |  |
| A363 | T96 (P12) |  |  |  |  |  |  |
| T366 | T96 (P12) | I97 (P12) |  |  |  |  |  |
| L370 | I97 (P12) |  |  |  |  |  |  |
| Q397 | V106 (P12) | T108 (P12) | L109 (P12) |  |  |  |  |
| P398 | L109 (P12) |  |  |  |  |  |  |
| N400 | Q105 (P12) | V106 (P12) |  |  |  |  |  |
| F401 | L102 (P12) | V106 (P12) | L109 (P12) | L110 (P12) |  |  |  |
| L404 | 101 (P12) | L102 (P12) | Q105 (P12) | V106 (P12) |  |  |  |
| T405 | I97 (P12) | L102 (P12) |  |  |  |  |  |
| Q408 | I97 (P12) | S98 (P12) | R101 (P12) | L102 (P12) |  |  |  |
| G409 | I97 (P12) |  |  |  |  |  |  |
| Q412 | I97 (P12) | S98 (P12) |  |  |  |  |  |
| Y516 | F98 (P2) | S100 (P2) |  |  |  |  |  |
| E517 | F98 (P2) |  |  |  |  |  |  |
| V520 | F98 (P2) |  |  |  |  |  |  |
| D561 | K201 (P2) |  |  |  |  |  |  |
| P563 | F98 (P2) | R99 (P2) | S100 (P2) | 101 (P2) |  |  |  |
| S566 | 96 (P2) | K97 (P2) | F98 (P2) | R99 (P2) |  |  |  |
| Y567 | K97 (P2) | F98 (P2) |  |  |  |  |  |
| V664 | P16 (P10) | F18 (P10) |  |  |  |  |  |
| R672 | F18 (P10) |  |  |  |  |  |  |
| A673 | 19 (P10) | K19(P10) |  |  |  |  |  |
| Q676 | F18 (P10) | K19 (P10) | N21 (P10) |  |  |  |  |
| E677 | N21 (P10) |  |  |  |  |  |  |
| W680 | N21 (P10) |  |  |  |  |  |  |
| D681 | 205 (P10) |  |  |  |  |  |  |
| V684 | L23 (P10) | G203 (P10) | H204 (P10) | K205 (P10) |  |  |  |
| G685 | H204 (P10) | K205 (P10) |  |  |  |  |  |
| N687 | E92 (P10) | D95 (P10) | G203 (P10) | H204 (P10) |  |  |  |
| K688 | V208 (P2) | H204 (P10) |  |  |  |  |  |
| Y689 | K205 (P2) | V208 (P2) |  |  |  |  |  |
| P691 | 12 (P2) | N13 (P2) | E92 (P10) | D95 (P10) |  |  |  |
| Q692 | P11 (P2) | A12 (P2) |  |  |  |  |  |
| N695 | N10 (P2) | 86 (P10) | K87 (P10) | I97 (P10) |  |  |  |
| L696 | N10 (P2) |  |  |  |  |  |  |
| I698 | L23 (P10) | I97 (P10) | G203 (P10) |  |  |  |  |
| M699 | P7 (P2) | I106 (P10) |  |  |  |  |  |
| W701 | N21 (P10) |  |  |  |  |  |  |
| Y702 | R105 (P10) | I106 (P10) |  |  |  |  |  |
| E703 | I106 (P10) |  |  |  |  |  |  |
| V705 | F18 (P10) | A20 (P10) | A20 (P10) | N21 (P10) | 22 (P10) |  |  |
| N706 | A20 (P10) | N21 (P10) | K104 (P10) |  |  |  |  |
| S709 | F18 (P10) | K19 (P10) | A20 (P10) |  |  |  |  |

**SUPPLEMENTARY FIGURES**

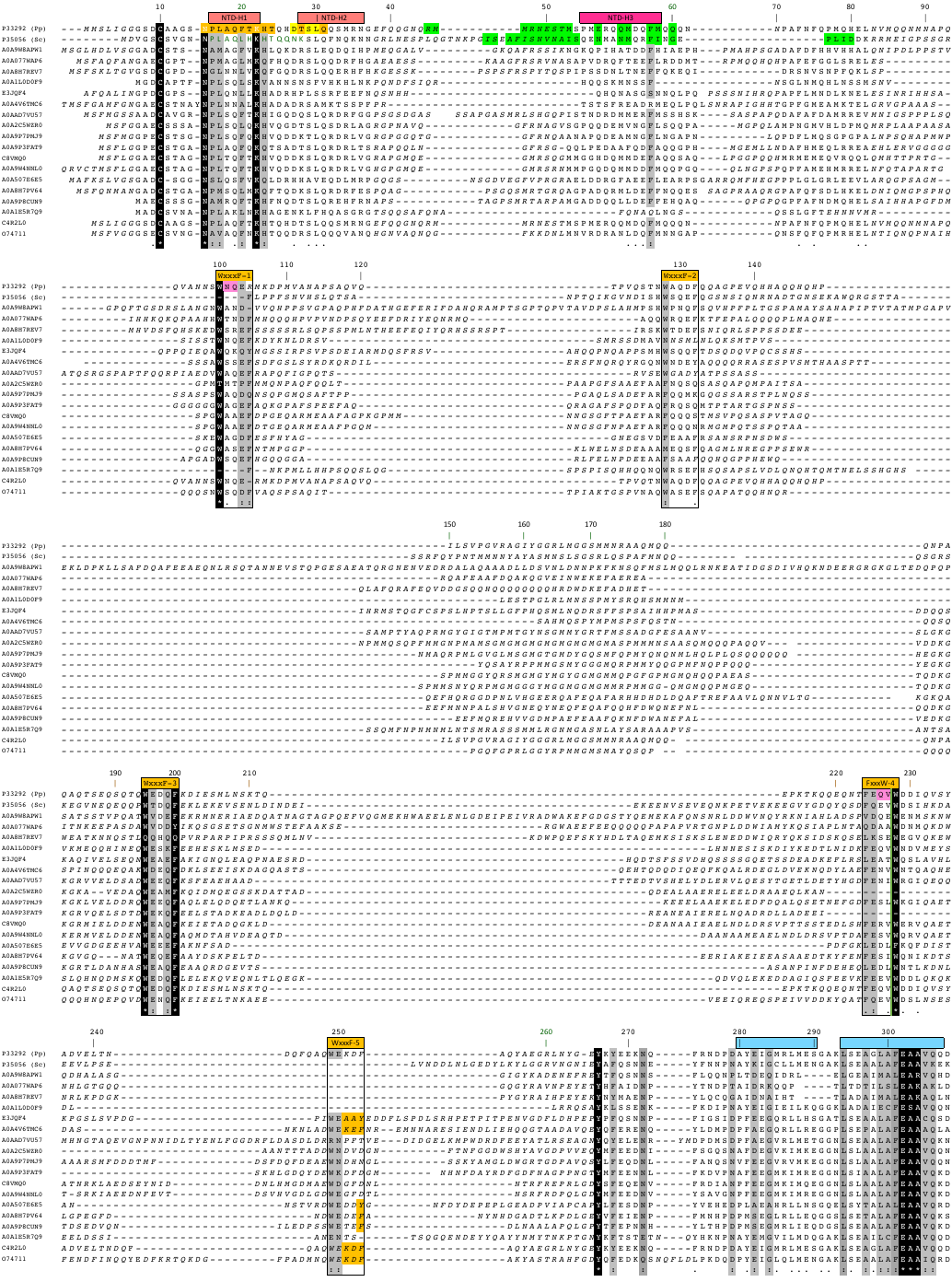

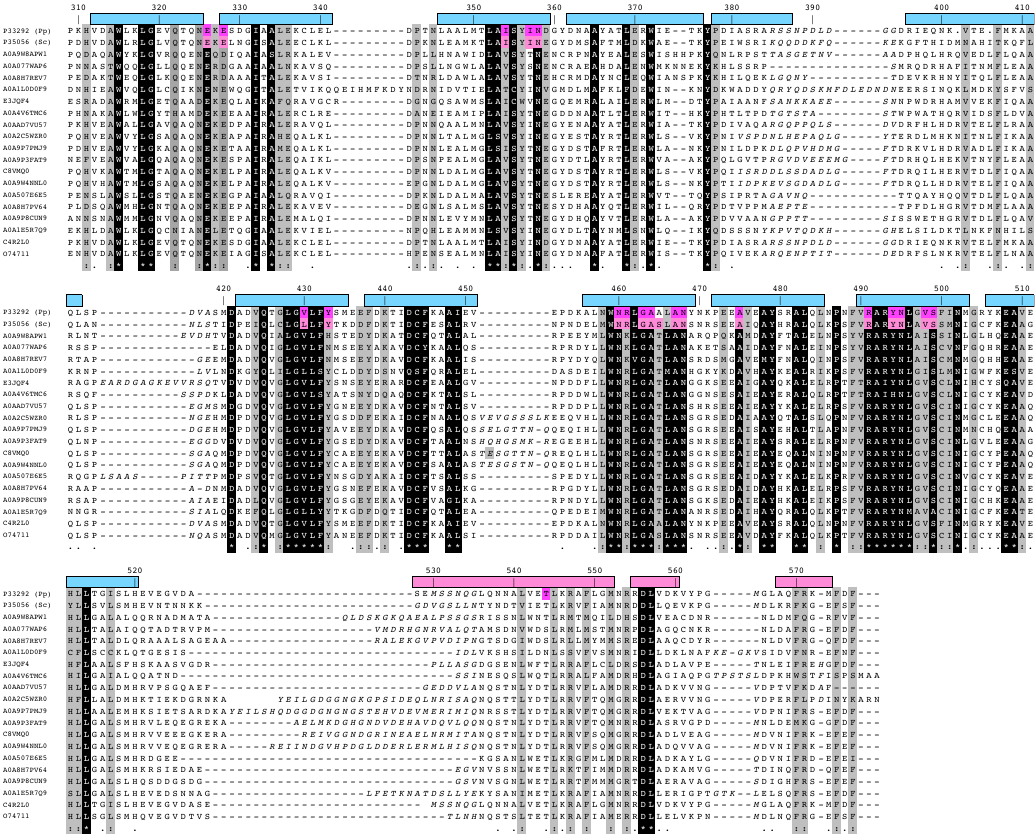

**Supplementary Figure 1. Multiple sequence alignment of representative fungi Pex5 sequences.** UNIPROT IDs are to the left. Unreliable parts of the alignment are in italics and have been condensed in parts for simplification. The Pex5 sequences from *P. pastoris* and *S. cerevisiae* are shown in the first two top lines. The numbers for every tenth residue correspond to the *P. pastoris* Pex5 sequence. Secondary structural elements are indicated above the alignment, adapting the color scheme used in the main figures. Experimentally determined and predicted interaction sites with Pex8, Pex2, Pex10, and Pex12 are shown in complementary colors of the respective protein ligand. Invariant and conserved residue positions are highlighted in black and grey, respectively. The bipartite structural organization of Pex5, which can be divided into a mostly unfolded NTD and a folded CTD, is directly reflected in the multiple sequence alignment. Whereas conserved sequence stretches in the Pex5 NTD are confined to short linear motifs close to the N-terminal cysteine ubiquitination site and an array of degenerative WxxxF motifs, the overall level of sequence conservation in the Pex5 CTD is considerably higher than in the Pex5 NTD. The alignment is based on BLAST searches of P33292 (Pp) against UNIPROT (Release 2025_01). Seed sequences sharing at least 50% identity with other cluster sequences (UNIREF50) are shown. Source organisms: P33292 (Pp), *Pichia pastoris*; P35056 (Sc), *Saccharomyces cerevisiae*; A0A9W8APW1, *Dispira parvispora*; A0A077WAP6, *Lichtheimia ramose*; A0A8H7REV, *Mucor plumbeus*; A0A1L0D0F9, *Hanseniaspora guilliermondii*; E3JQF4, *Puccinia graminis*; A0A4V6TMC6, *Wallemia hederae*; A0AAD7VU57, *Lipomyces tetrasporus*; A0A2C5WZR0, *Ceratocystis fimbriata*; A0A9P7PMJ9, *Claviceps sp*; A0A9P3FAT, i; C8VMQ0, *Emericella nidulans*, A0A9W4NNL0, *Penicillium salami*; A0A507E6E5, *Powellomyces hirtus*; A0A8H7PV64, *Absidia glauca*; A0A9P8CUN, *Mortierella alpina*; A0A1E5R7Q9, *Hanseniaspora osmophila*; C4R2L0, *Komagataella phaffii*; O74711, *Candida albicans*.

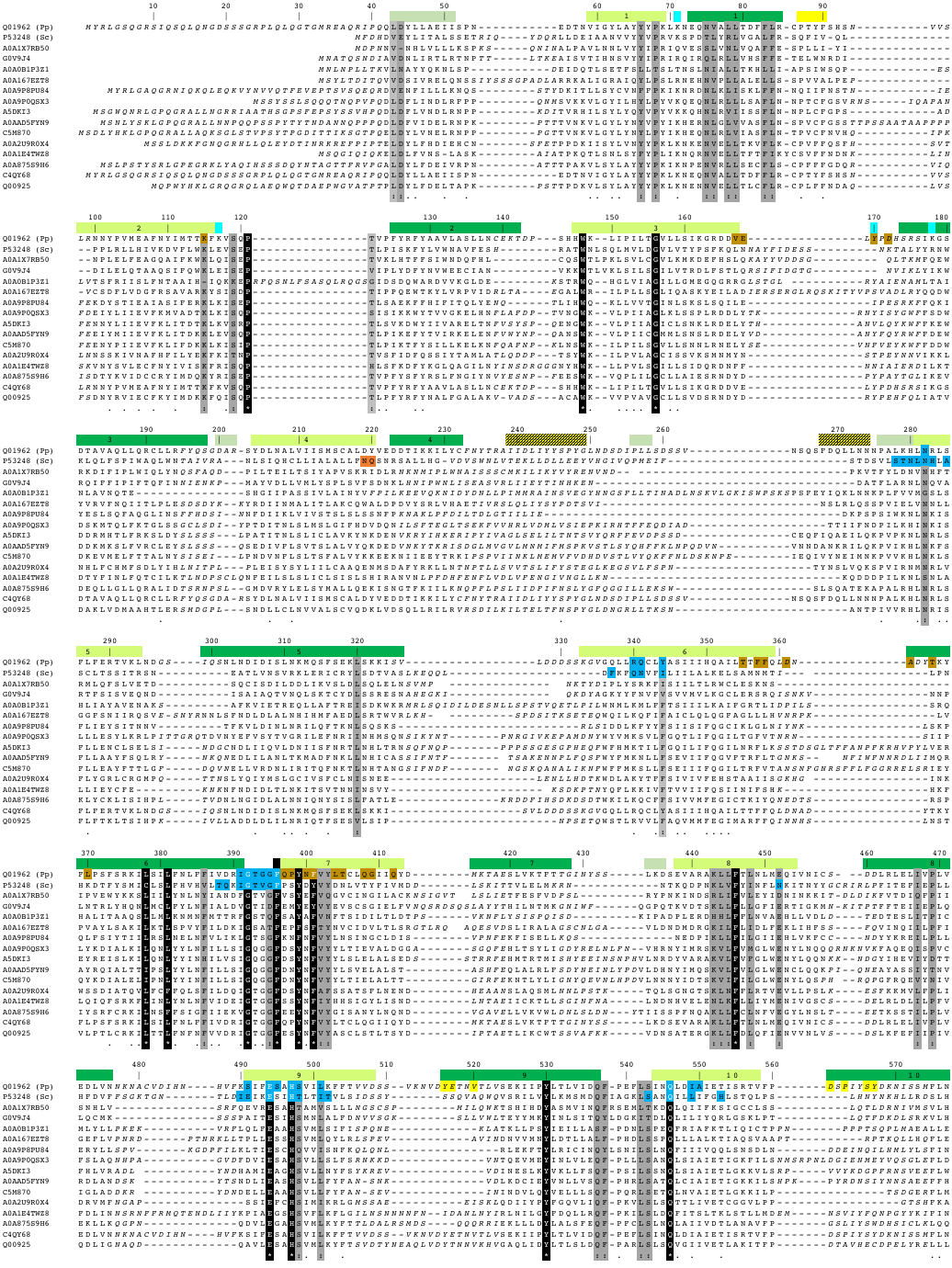

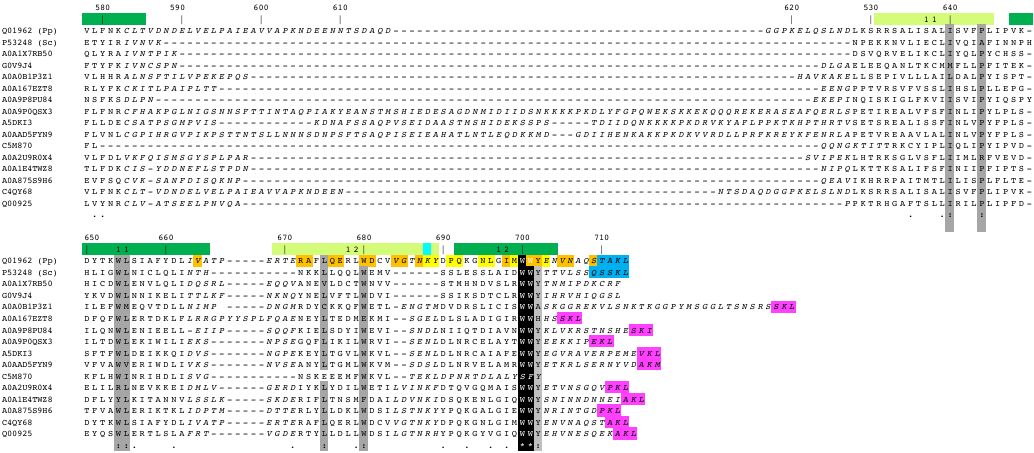

**Supplementary Figure 2. Multiple sequence alignment of representative fungi Pex8 sequences.** UNIPROT IDs are to the left. Unreliable parts of the alignment are in italics and have been condensed in parts for simplification. The Pex8 sequences from *P. pastoris* and *S. cerevisiae* are shown in the first two top lines. The numbers for every tenth residue correspond to the *P. pastoris* Pex8 sequence. Secondary structural elements are indicated above the alignment, adapting the color scheme used in the main figures. Experimentally determined and predicted interaction sites with Pex5, Pex2, Pex10, Pex12 are shown in complementary colors of the respective protein ligand. Invariant and conserved residue positions are highlighted in black and grey, respectively. The C-terminal PTS1 motif is not universally conserved among the sequences shown in this alignment. The alignment is based on BLAST searches of Q01962 (Pp) against UNIPROT (Release 2025_01). Seed sequences sharing at least 50% identity with other cluster sequences (UNIREF50) are shown. Source organisms: Q01962 (Pp), *Pichia pastoris*; P53248 (Sc), *Saccharomyces cerevisiae*; A0A1X7RB50, *Maudiozyma saulgeensis*; G0V9J4, *Naumovozyma castellii*; A0A0B1P3Z1, *Uncinula necator*; A0A167EZT, *Sugiyamaella lignohabitans*; A0A9P8PU84, *Wickerhamomyces mucosus*; A0A9P0QSX, *Candida railenensis*; A5DKI3, *Meyerozyma guilliermondii*; A0AAD5FYN9, *Candida theae*; C5M870, *Candida tropicalis*; A0A2U9R0X4, *Pichia kudriavzevii*, A0A1E4TWZ8, *Pachysolen tannophilus*; A0A875S9H6, *Eeniella nana*; C4QY68, *Komagataella phaffii*, Q00925, *Pichia angusta*.

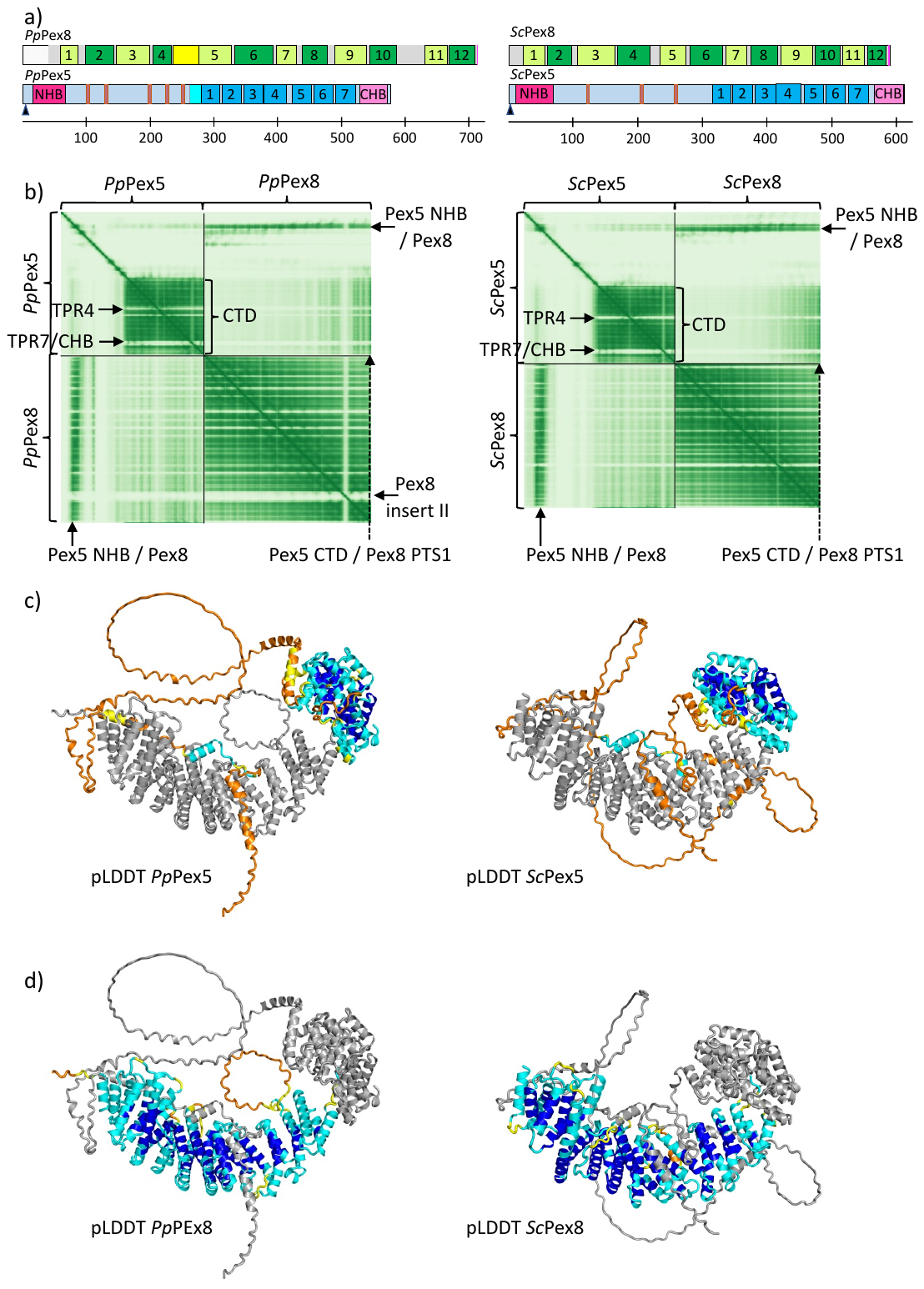

**Supplementary Figure 3. AF3 models of the Pex5/Pex8 complexes from *P. pastoris* (left) and *S. cerevisiae* (right). a,** scheme of the structural/functional organization of the Pex5 and Pex8 sequences from *P. pastoris* and *S. cerevisiae*. Colors are adapted from those in the main figures; **b,** PAE plots of the AF3-predicted Pex5/Pex8 models from *P. pastoris* (left panel) and *S. cerevisiae* (right panel). Several low-scoring (faint colors) and high-scoring (strong green) interaction sites are indicated and labeled. The plots are proportional to sequence length; **c,** AF3 Pex5/Pex8 models from *P. pastoris* (left panel) and *S. cerevisiae* (right panel), in which Pex5 is colored by AF3 pLDDT scores; **d,** AF3 Pex5/Pex8 models from *P. pastoris* (left panel) and *S. cerevisiae* (right panel), in which Pex8 is colored by AF3 pLDDT scores. Whereas most of the Pex8 sequence is predicted with high local confidence scores, in Pex5 they are confined to its folded CTD and a short sequence stretch within the NHB segment of the Pex5 NTD (for further details see **Figure 5**).

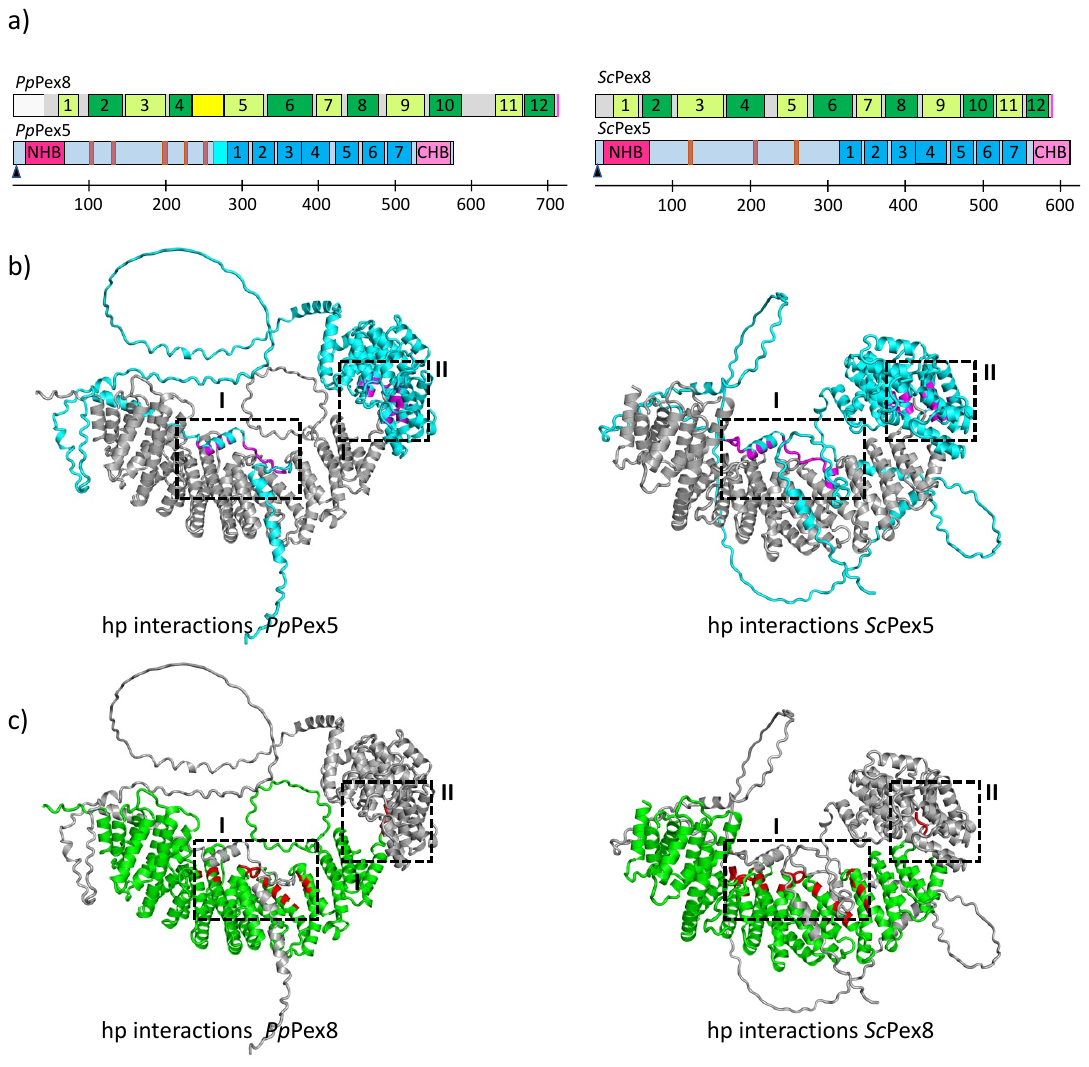

**Supplementary Figure 4. AF3 models of the Pex5/Pex8 complexes from *P. pastoris* (left) and *S. cerevisiae* (right), continued from Supplementary Figure 3**. **a,** identical with panel a of **Supplementary Figure 3; b,** AF3 Pex5/Pex8 models from *P. pastoris* (left panel) and *S. cerevisiae* (right panel), in which Pex5 is colored in cyan. High-confidence interaction sites with Pex8 **(Supplementary Table 2)** are colored in magenta**; c,** AF3 Pex5/Pex8 models from *P. pastoris* (left panel) and *S. cerevisiae* (right panel), in which Pex8 is colored in green. High-confidence interaction sites with Pex5 **(Supplementary Table 2)** are colored in red.

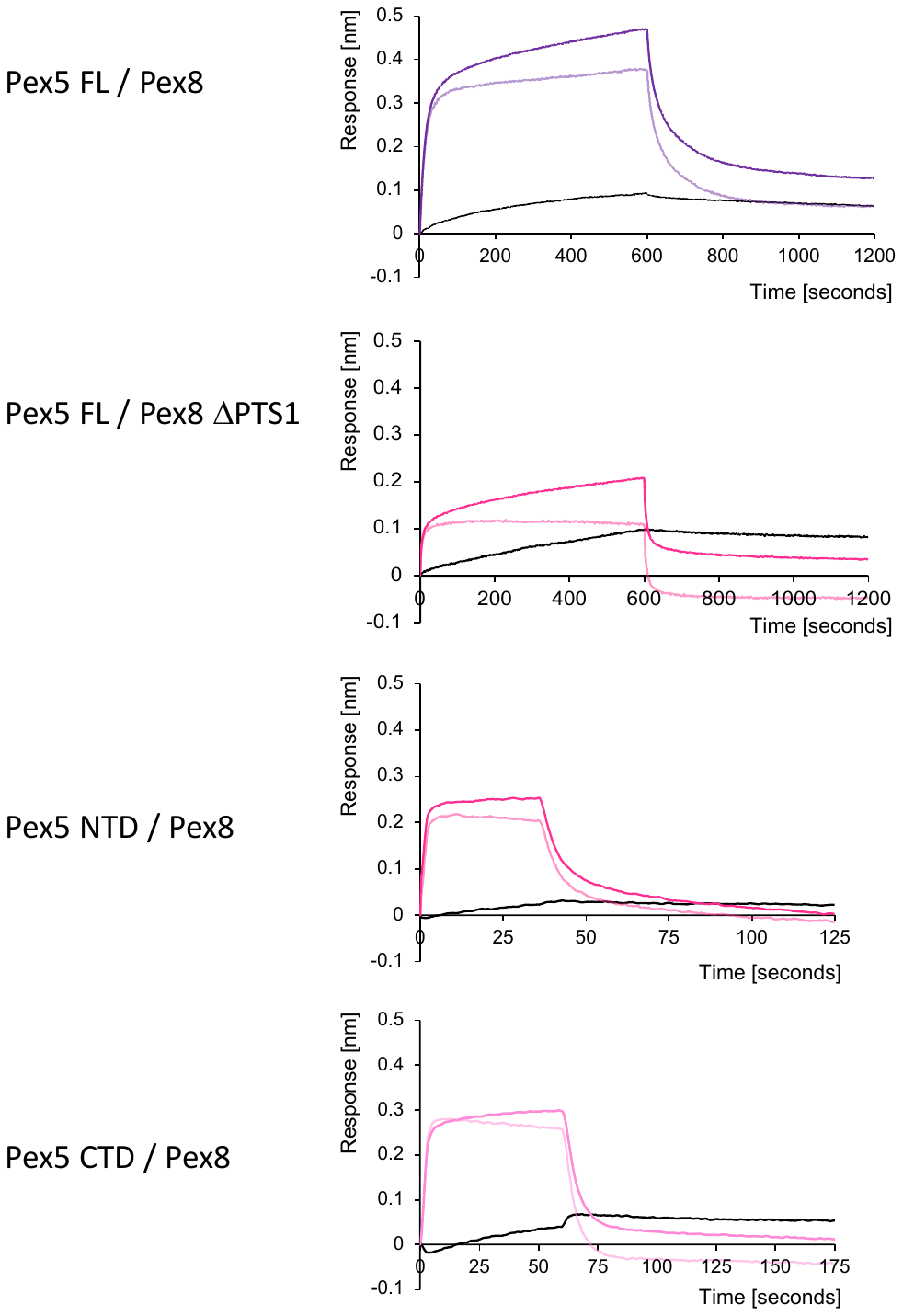

**Supplementary Figure 5. Representative BLI Octet curves of Pex5 FL, NTD and CTD interacting with Pex8 WT and Pex8 ΔPTS1.** For each interaction, the association/dissociation curves using the original data (saturated) and background-subtracted data (half saturated), as well as background curves (black) are shown. Color codes of sample curves are as in **Figure 2 b-c.**

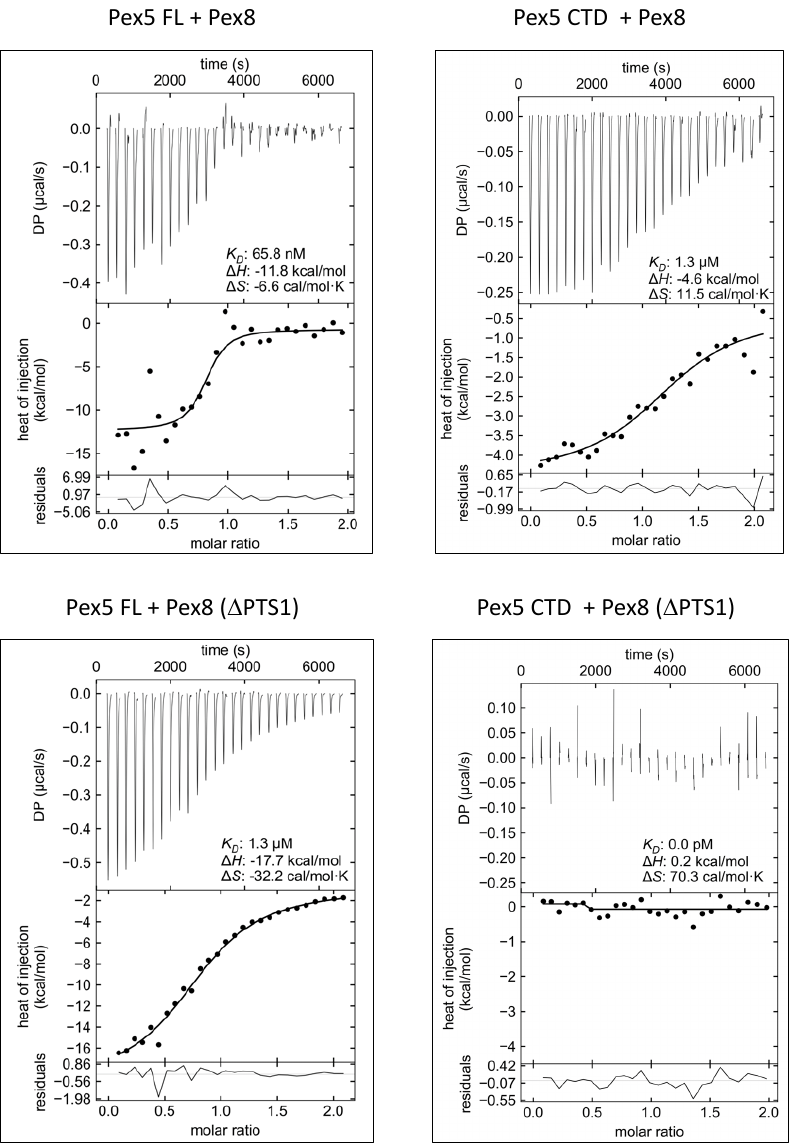

**Supplementary Figure 6.** **Representative ITC curves of complex formation by different Pex5 and Pex8 constructs.** The plots show the reconstructed thermograms based on singular value decomposition, indicated by the differential power (DP) versus time (s) (upper panel), heat of injection (kcal/mol) (central panel), and residual plots as function of molar ratios (lower panel). The calculated dissociation constants K_D_, enthalpy terms (ΔH) and entropy terms (ΔS) are presented for each curve.

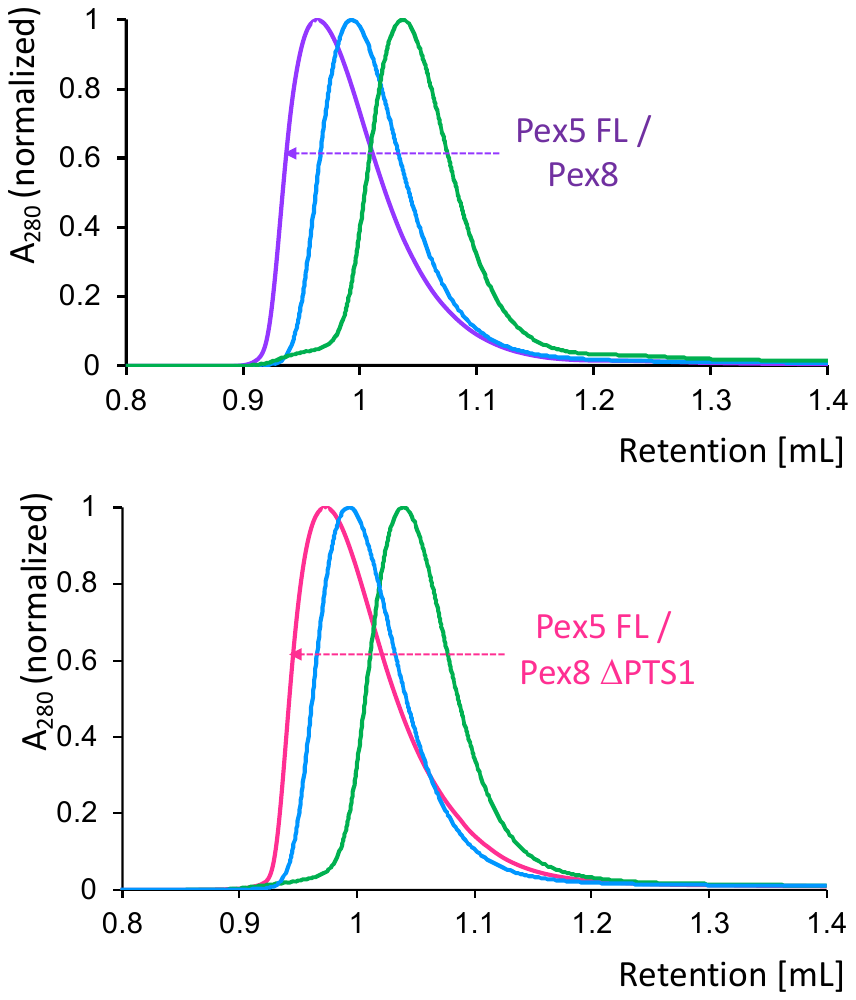
**Supplementary Figure 7. Normalized SEC profiles of Pex5 FL in complex with WT Pex8 and Pex8 ΔPTS1.** Color codes are as in **Figure 2.** The SEC profiles of the separate Pex5 and Pex8 constructs are in marine and green, respectively. The SEC profiles in complex with WT Pex8 and Pex8 ΔPTS1 are indistinguishable in terms of the observed retention volume. The data were obtained from an analytical S75 3.2/300 Column (Cytiva).

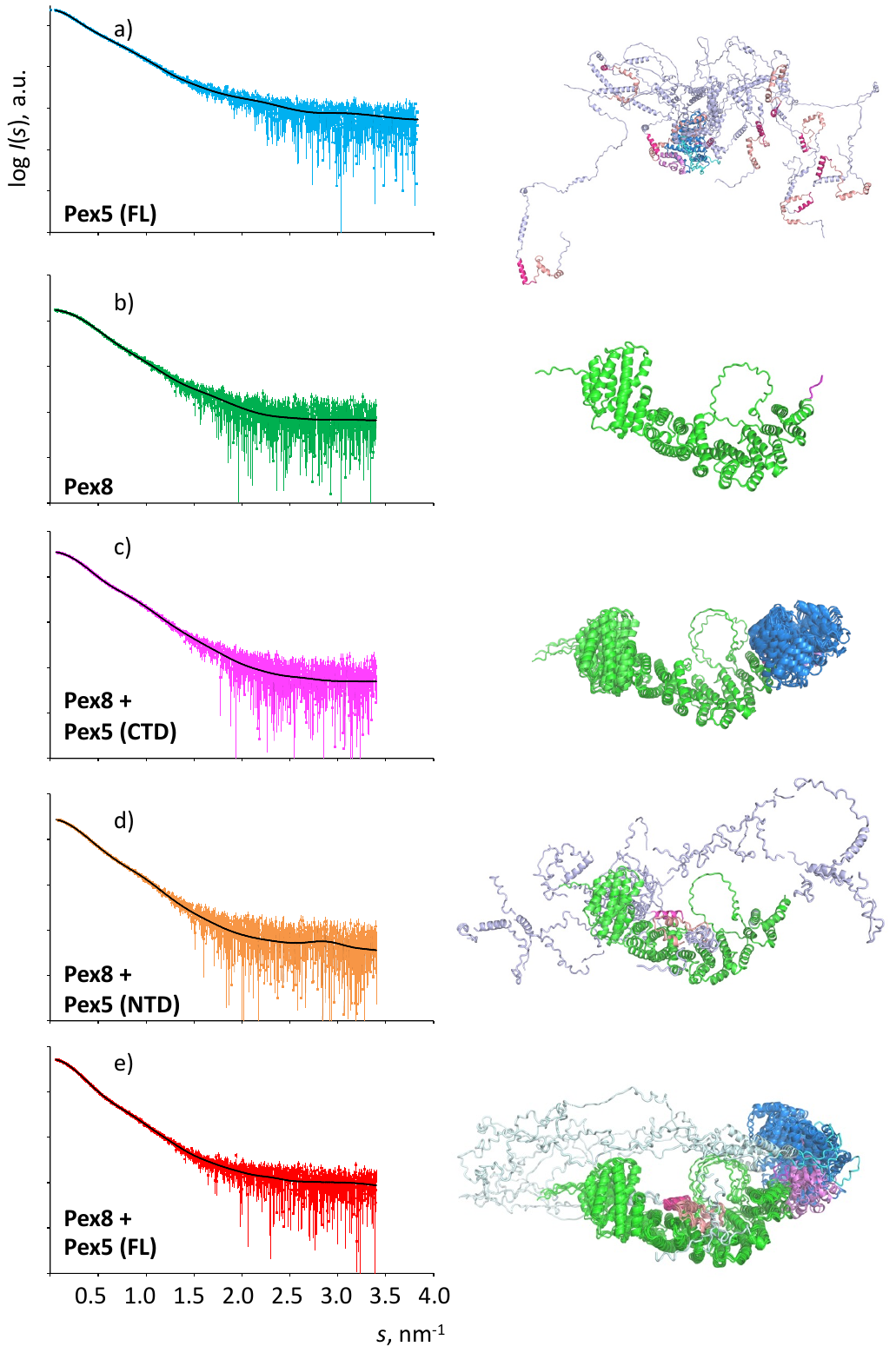

**Supplementary Figure 8. SAXS curves of different Pex5/Pex8 complexes (left) and complete model ensembles (right).** The SAXS curves are shown as logarithmic I(s) plots in arbitrary units *versus* reciprocal resolution (nm^-1^). All plots are at the same resolution scale. Colors are as in **Figure 3a**. The complete model ensembles are shown in cartoon presentation, using color codes adopted from **Figure 3b-f.** The data indicate limited conformational variability of the Pex5 CTD when bound to the Pex8 PTS1 motif. The models also support a well-defined Pex5 NHB/Pex8 binding site. Conformational diversity of the remaining Pex5 sequence is restricted in the presence of the second Pex5 CTD/Pex8 PTS1 interaction site. In the absence of Pex8 (or other ligands), there is no restriction in Pex5 conformational diversity, except for the folded CTD.

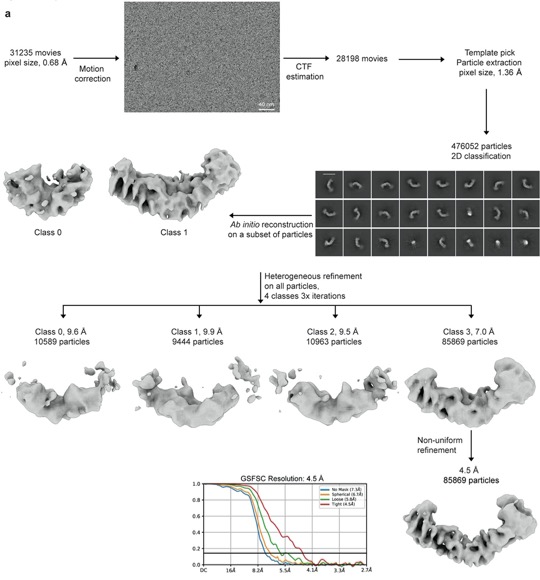

**Supplementary Figure 9. Flowcharts** **for cryo-EM analysis and integrative modeling of the Pex5/Pex8 complex. a,** cryo-EM data processing workflow applied to 31,235 movies using CryoSPARC **[1],** resulting in a 4.5 Å resolution density map of the Pex5/Pex8 complex **(Figure 4c-d).** Data collection and refinement statistics are listed in **Supplementary Table 3**.

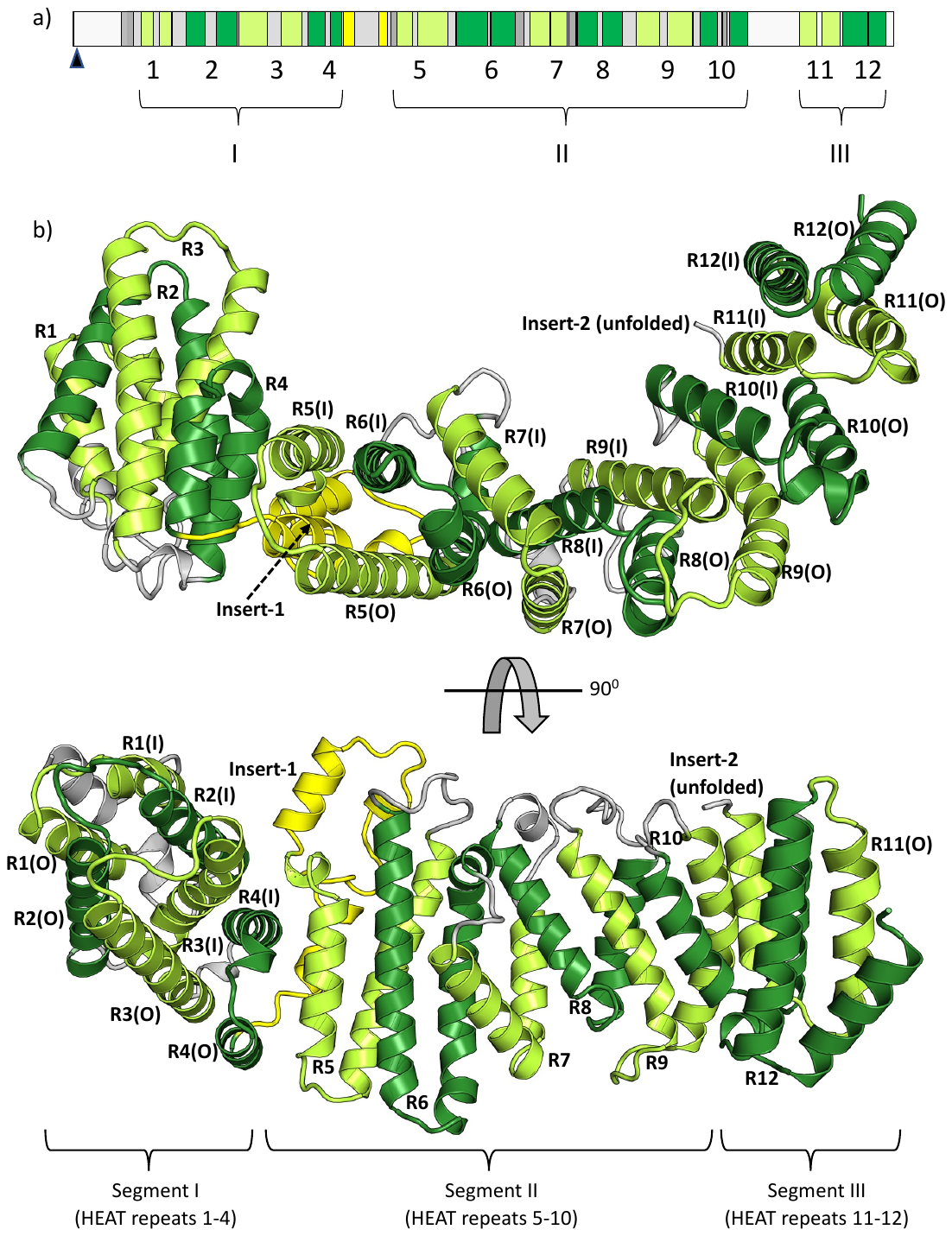

**Supplementary Figure 10. Crystal structure of Pex8. A,** scheme of the structural/functional organization of the *P. pastoris (Pp)* Pex8 sequences. In this scheme, individual HEAT repeats and insert-1 helices are shown separately, in colors adapted from the main figures (limon, green, yellow). Other helices are in dark grey, and the remaining visible regions of the Pex8 sequence are in light grey. HEAT repeats and domain segments are numbered and labeled; **b,** cartoon representation of the Pex8 crystal structure, in two different orientations, rotated by 90 degrees around a horizontal axis.

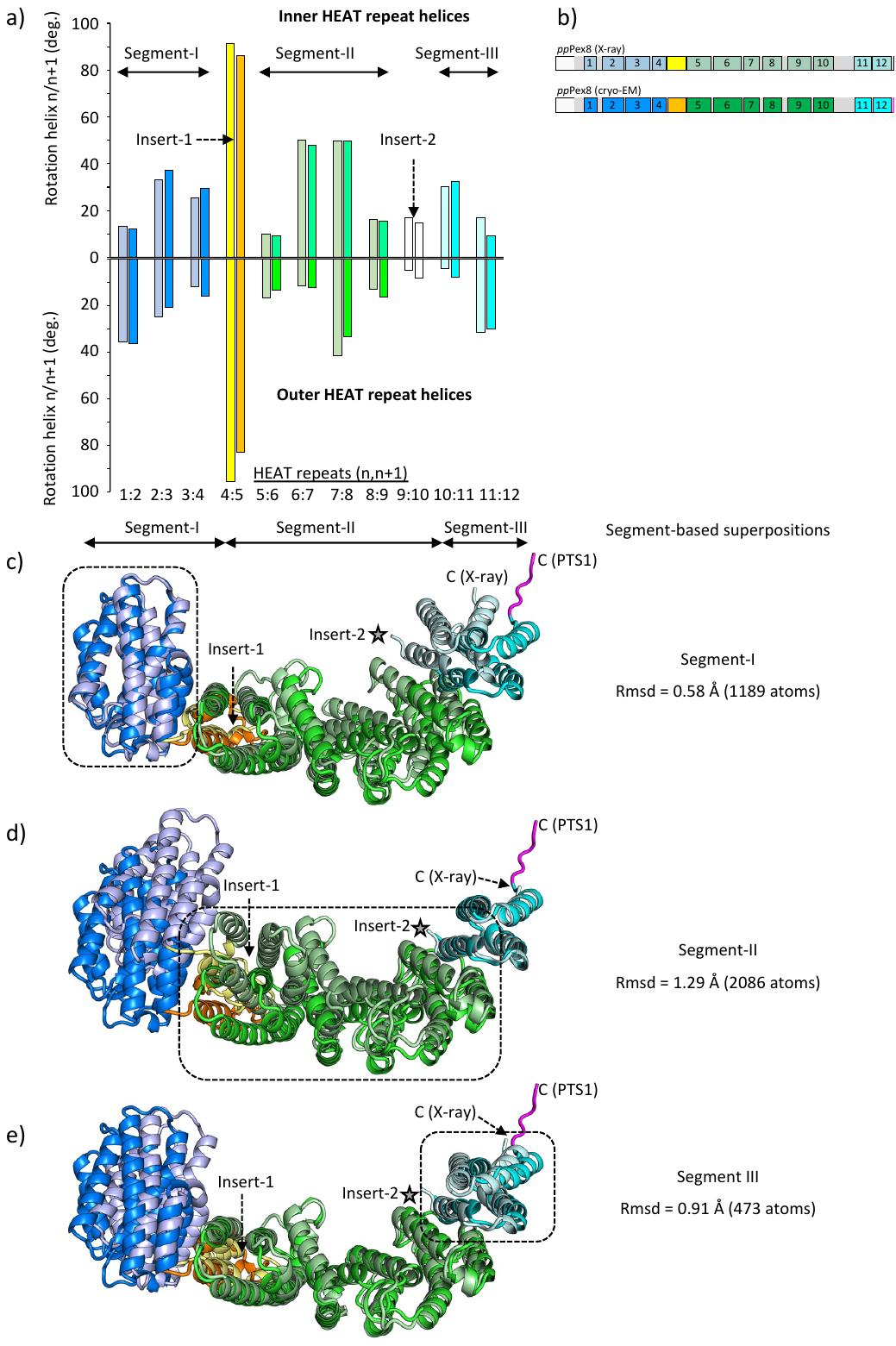

**Supplementary Figure 11. Comparison of the crystal and cryo-EM Pex8 structures. a,** angles between main axes of neighboring HEAT repeat helices, taken from the respective crystal structure (faint colors) and cryo-EM structure (bold colors). Color codes: segment I HEAT repeat arrangements, faint and strong blue; HEAT repeats 4/5 arrangement, bridging segments I and II and covering the structured Insert-1 domain, yellow/orange; segment II HEAT arrangements, faint and strong green; HEAT repeats 9/10 arrangement, bridging segments II and III and covering the unstructured insert-2, uncolored; segment III HEAT repeat arrangements, faint and strong cyan. Upper histogram, inner HEAT repeat helices; lower histogram, outer HEAT repeat helices. For further details, see **Supplementary Table 5; b,** scheme of the structural/functional organization of the *P. pastoris (Pp)* Pex8 sequences, highlighting structural elements of the Pex8 crystal structure (**Supplementary Figure 10**) and the Pex8 structure, taken from the Pex5/Pex8 complex determined by cryo-EM **(Figure 4e).** The color scheme is as in panel a; **c,** superpositions of the Pex8 crystal structure onto the Pex8 structure taken from the Pex5/Pex8 complex determined by cryo-EM, using segments I, II and III as base. These superpositions reveal substantial deviations, exceeding 10 Å distances especially at the Pex8 poles, opposite to the superimposed segments. Colors are as in panels a and b.

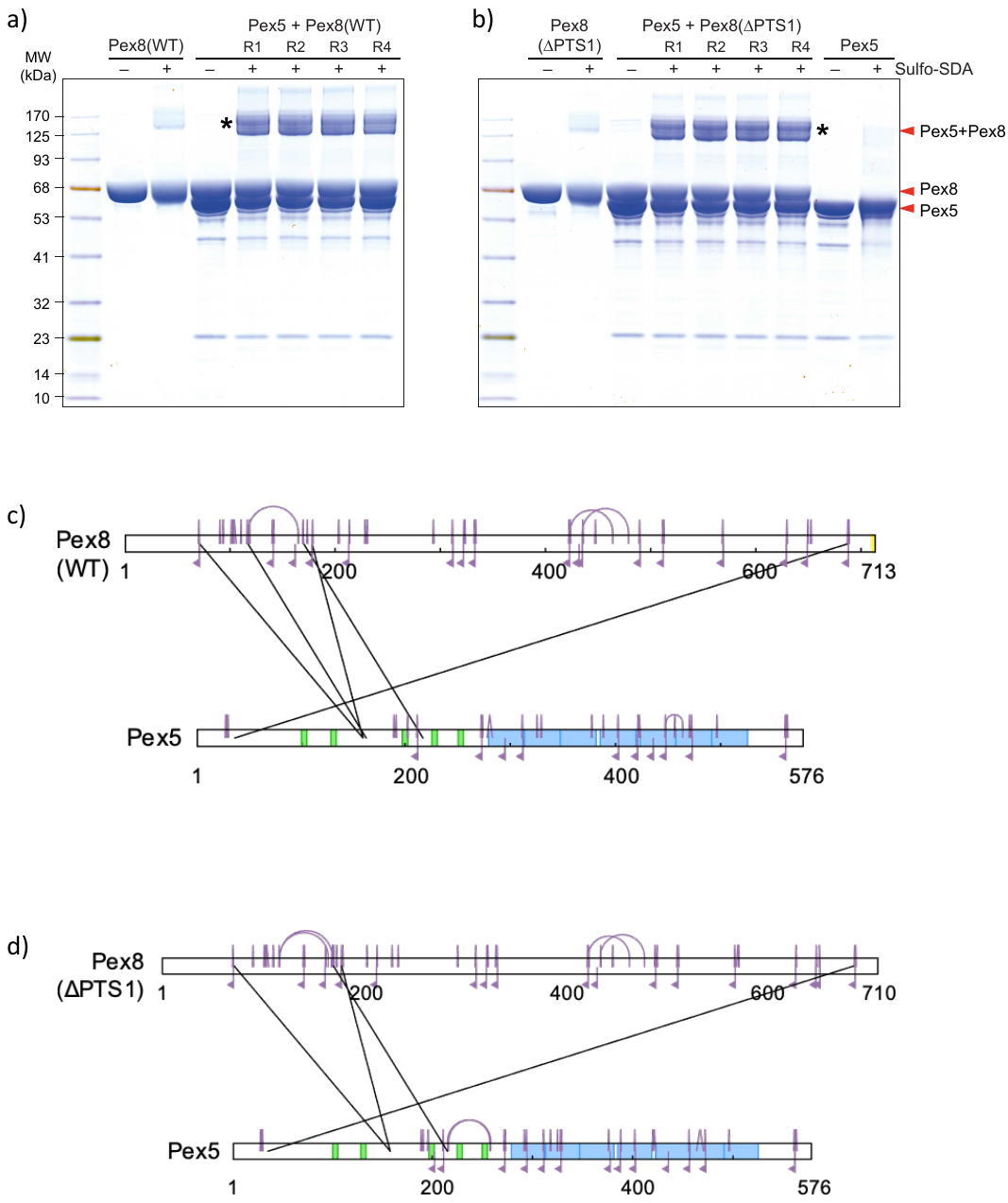

**Supplementary Figure 12. Crosslinking mass spectrometry (XL-MS) analysis of the Pex5/Pex8 complex. a,b** Coomassie-stained gel showing Pex5, Pex8 and Pex5/Pex8 complex samples after completing the crosslinking reaction. Bands corresponding to Pex5/Pex8 complexes indicated by asterisks (*) were excised for mass spectrometry analysis. **c,** schematic diagram showing the crosslinks identified by mass spectrometry within and between the Pex8 (WT, ΔPTS1) and Pex5 components of the Pex5/Pex8 complex. Colors: Intramolecular crosslinks and monolinks, purple; intermolecular crosslinks, black.

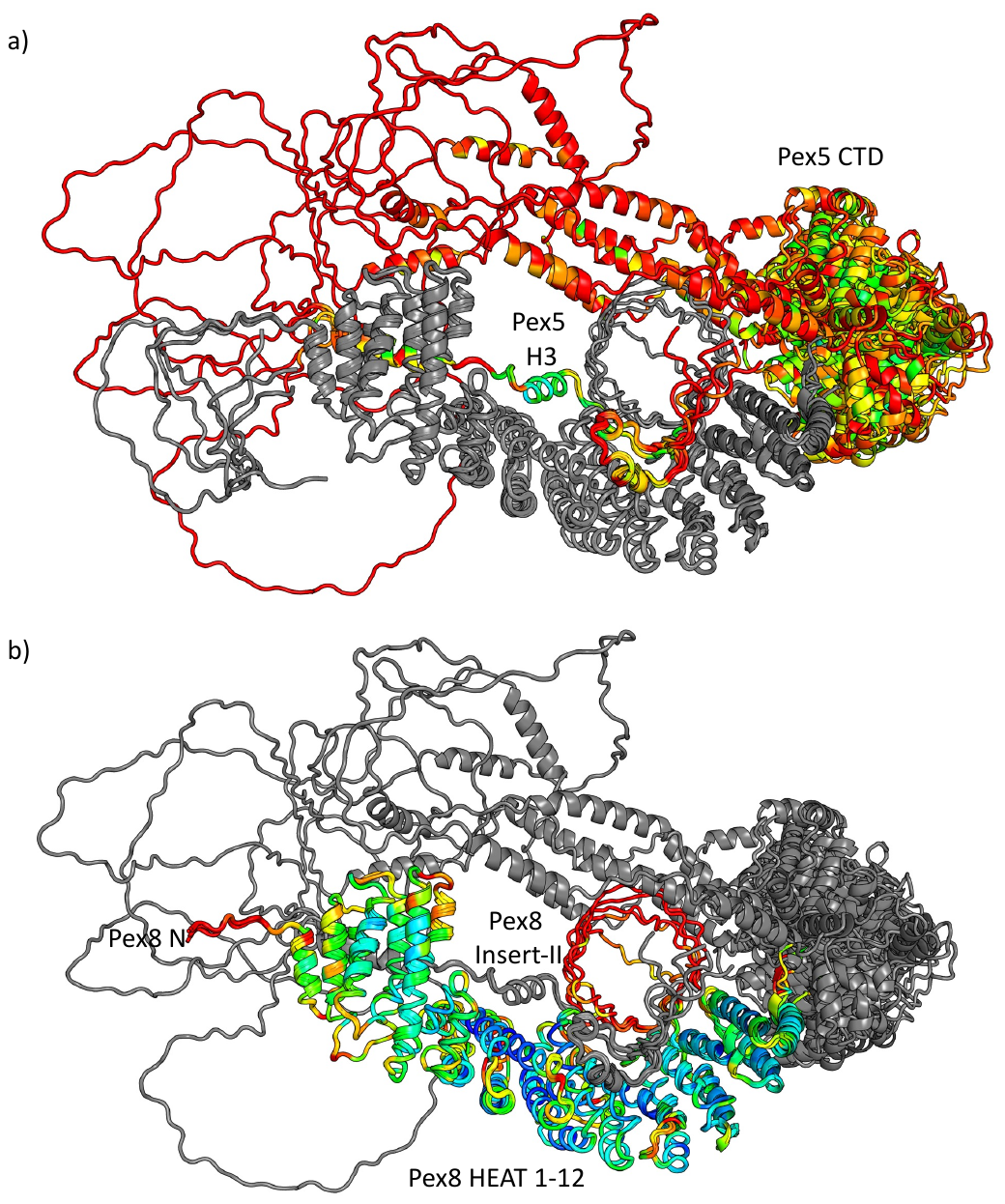

**Supplementary Figure 13. EM-data based Pex5/Pex8 integrative model ensemble. a,** ensemble model highlighting Pex5 by B-factor range colors; **b,** same ensemble model highlighting Pex8 by B-factor range colors. Sequence segments, for which no density could be detected or for which the density was of insufficient resolution to allow any further structural interpretation are generally colored in orange to red.

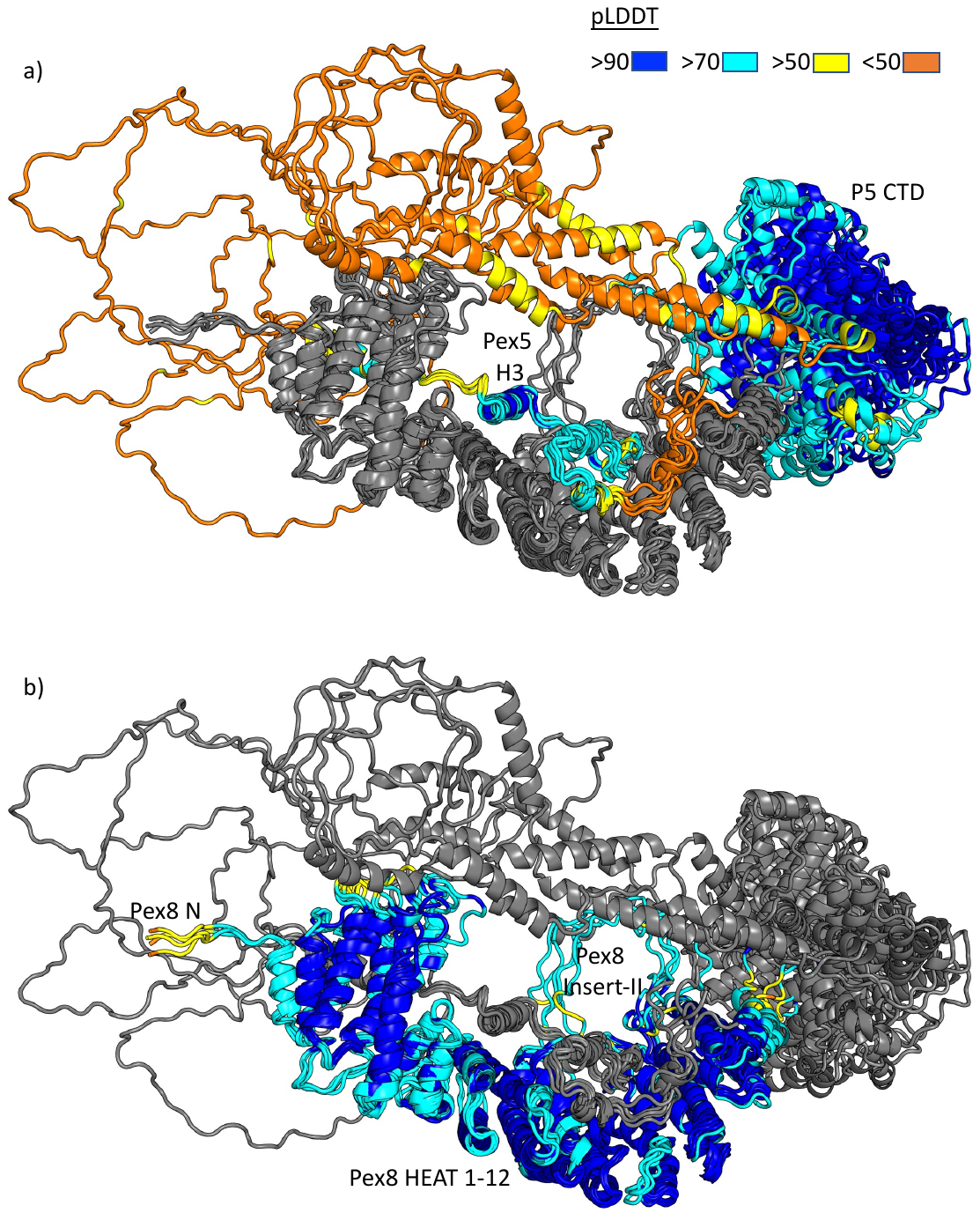

**Supplementary Figure 14. XL-MS data based Pex5/Pex8 integrative model ensemble. a,** ensemble model highlighting Pex5 by pLDDT color range; **b,** same ensemble model highlighting Pex8 by pLDDT color range. Sequence segments with low pLDDT scores are colored in orange to yellow.

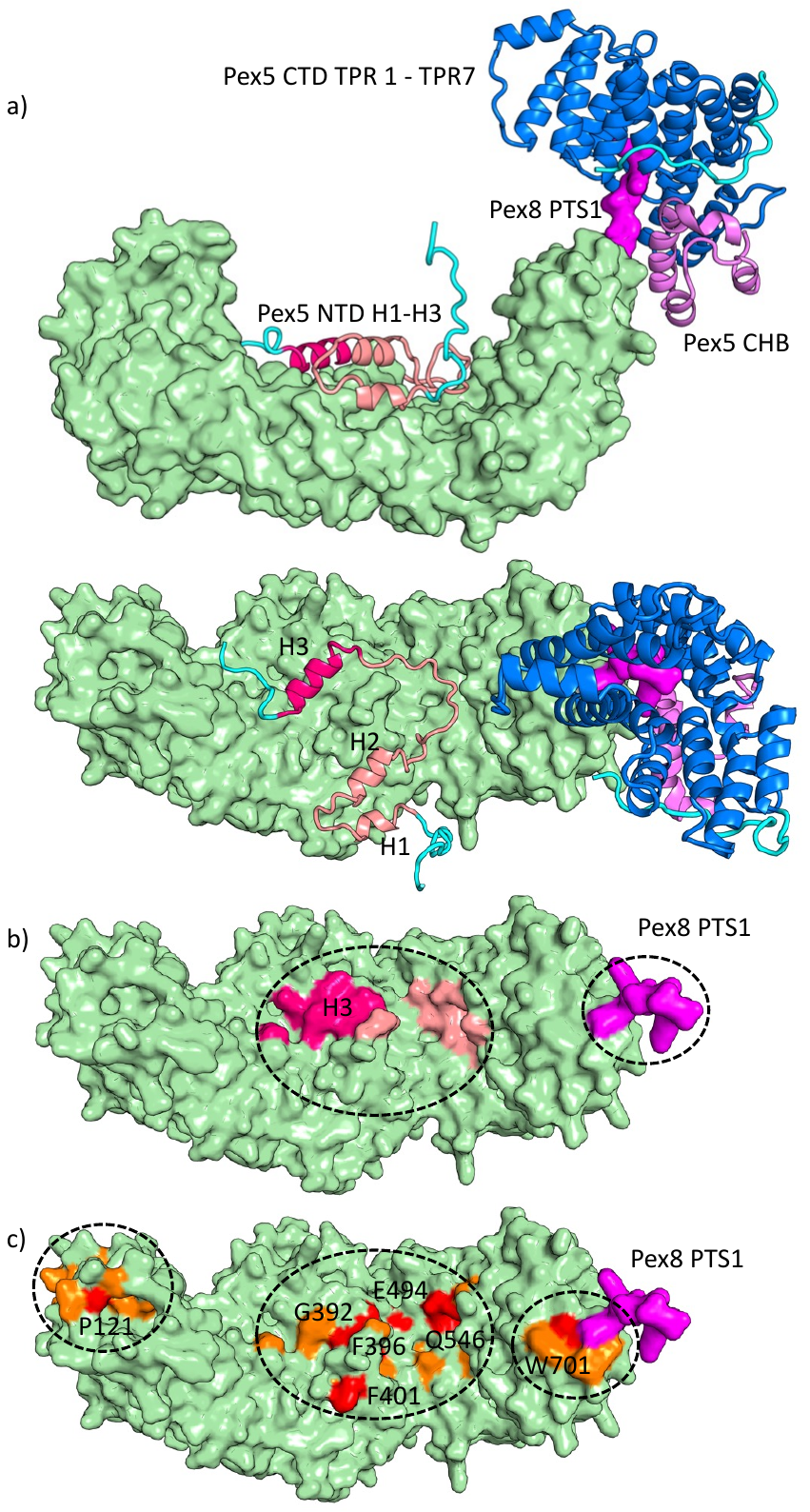
**Supplementary Figure 15. Pex8 surface highlighting the Pex5 NHB/Pex8 and Pex5 CTD/Pex8 PTS1 interaction sites. a,** complex of Pex8 with Pex5, in two different orientations rotated by approximately 90 degrees around a horizontal axis. For clarity, only the structure of the Pex5 NHB and CTD is shown in cartoon presentation; **b,** Pex5 NHB interactions are mapped onto Pex8 surface in complementary colors, used for Pex5; **c,** invariant (red) and conserved (orange) surface patches, based on Pex8 multiple sequence alignment **(Supplementary Figure 2).** Unless specifically mentioned, other colors are adopted from the main figures. The two characterized Pex5 NHB/Pex8 and Pex5 CTD/Pex8 PTS1 interaction sites as well as the third Pex5/Pex8 close neighborhood sites are indicated by dashed ellipsoids, demonstrating that there is correlation with conserved residues on the Pex8 surface.

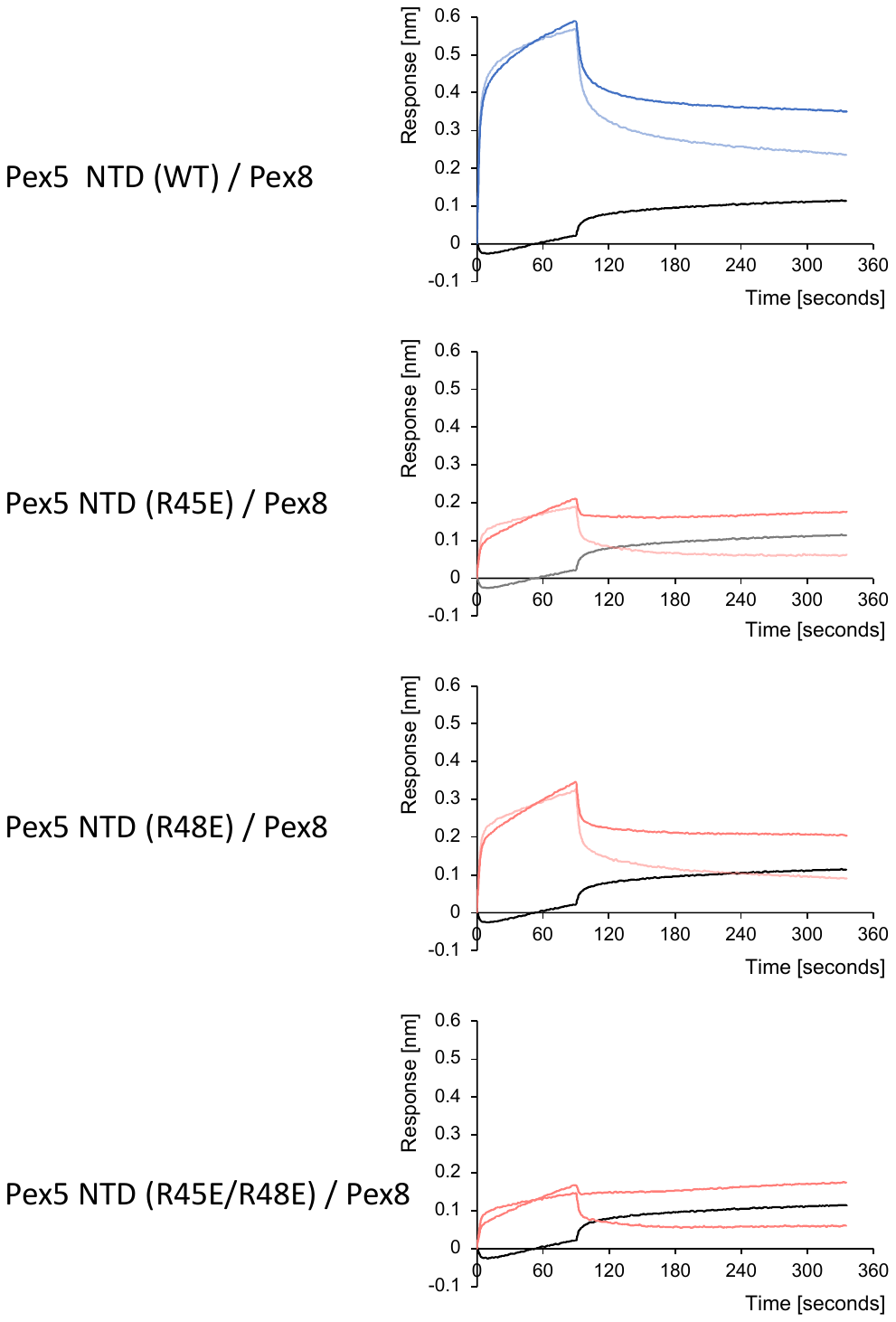

**Supplementary Figure 16. Representative BLI Octet curves of Pex5 NTD WT, R45E, R48E and R45E/R48E variants with Pex8 WT.** For each interaction, the association/dissociation curves using the original data (saturated) and background-subtracted data (half saturated), as well as background curves (black) are shown. Color codes of sample curves are as in **Figure 6c.**

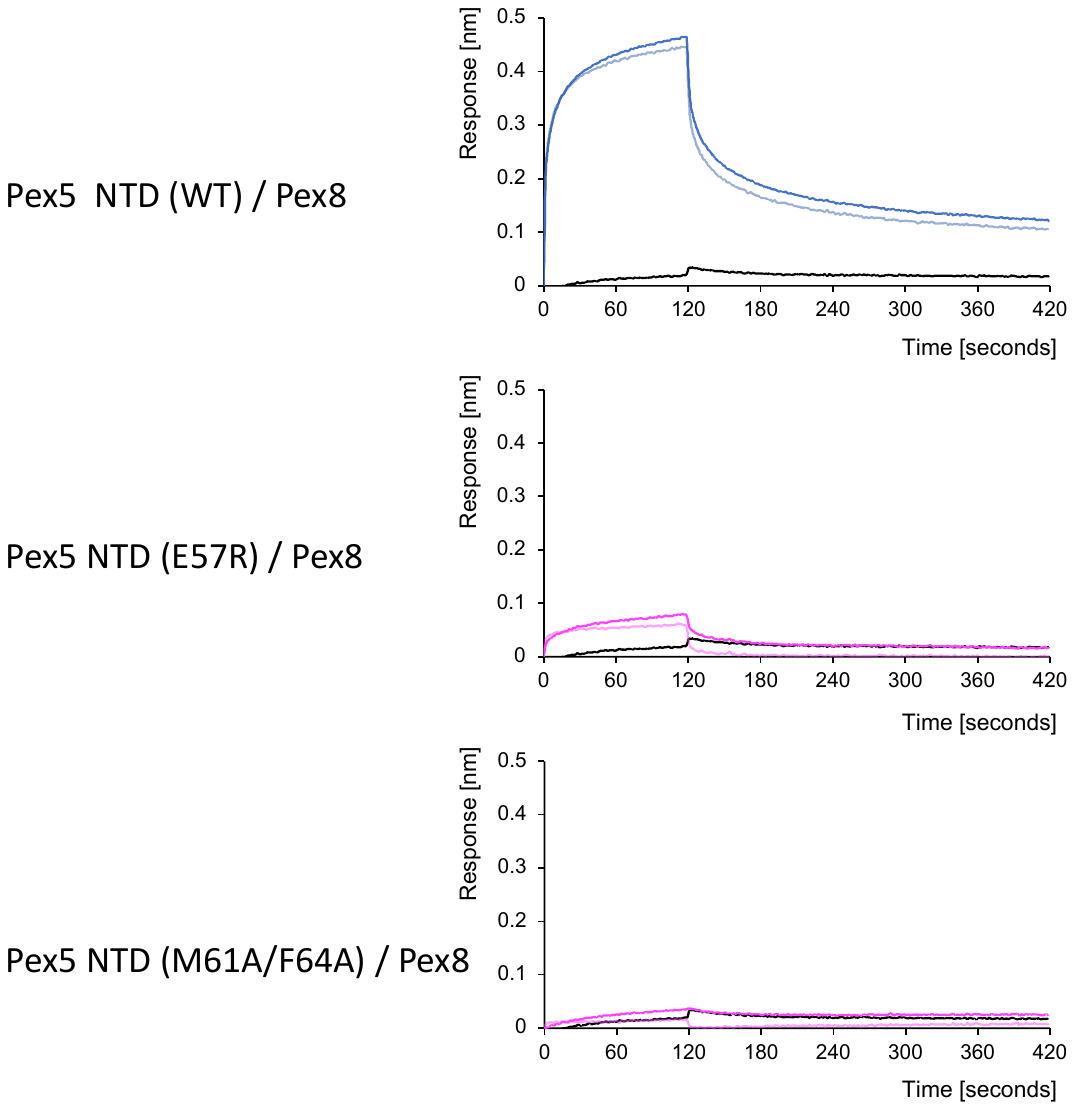

**Supplementary Figure 17. Representative BLI Octet curves of Pex5 NTD WT, E57R, and M61A/M64A variants with Pex8 WT.** For each interaction, the association/dissociation curves using the original data (saturated) and background-subtracted data (half saturated), as well as background curves (black) are shown. Color codes of sample curves are as in **Figure 6c.**

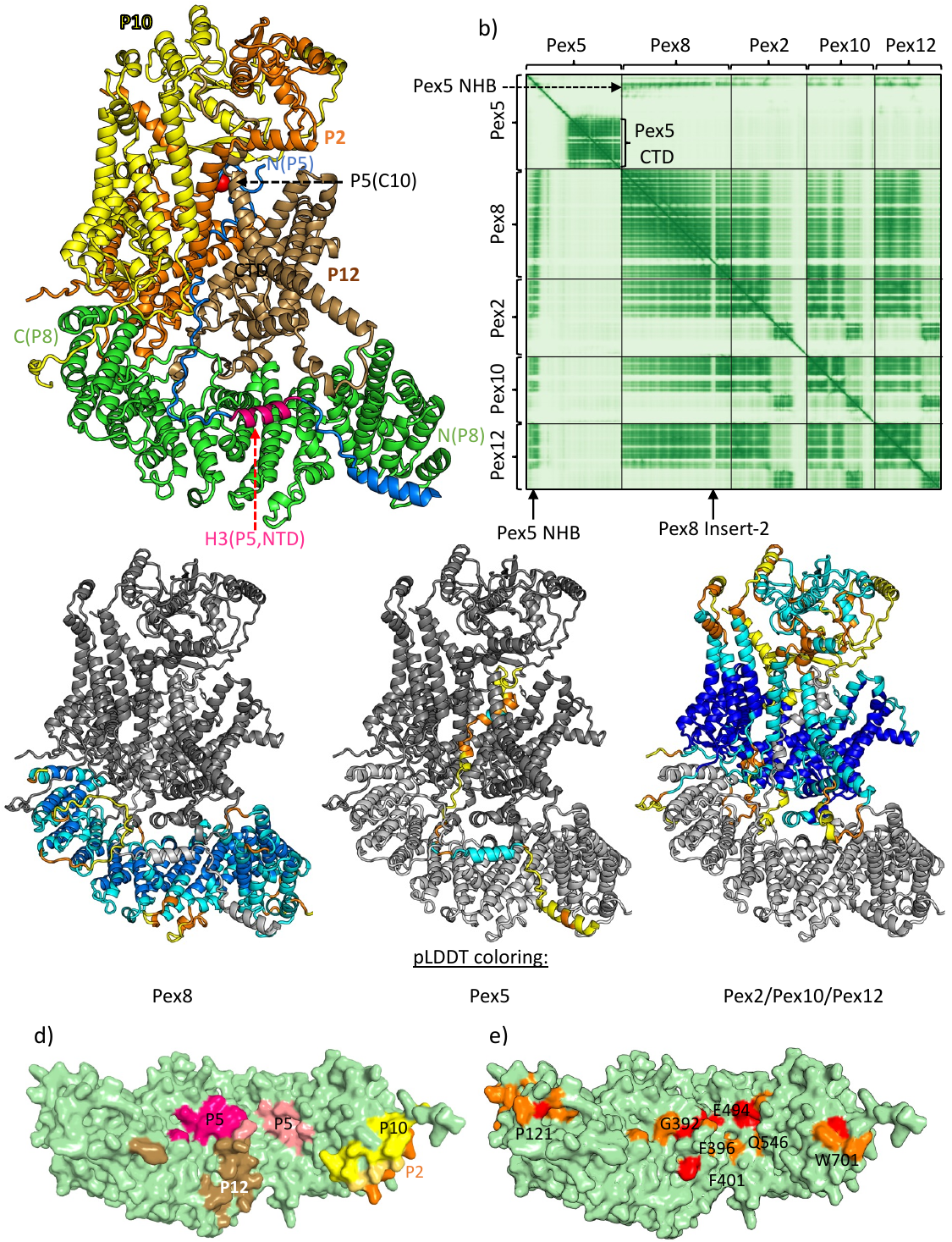
**Supplementary Figure 18. Details of the predicted Pex5/Pex8:Pex2/Pex10/Pex12 AF3 model. a,** cartoon representation of Pex5/Pex8:Pex2/Pex10/Pex12 model (*cf*. **Figure 7c**); **b,** PAE plot of the same model; **c,** Pex5/Pex8:Pex2/Pex10/Pex12 models with Pex8 (left), Pex5 (central) and Pex2/Pex10/Pex12 (right) highlighted in pLDDT score colors; **d,** surface representation of Pex8 with predicted Pex2/Pex10/Pex12 interaction sites in complementary colors added (*cf*. **Supplementary Figure 15b**); **e,** Pex8 surface representation highlighting invariant (red) and conserved (orange) surface patches, based on Pex8 multiple sequence alignment **(Supplementary Figures 2 and 15c).** Comparison reveals an assignment of the highly conserved WWY motif within HEAT repeat 12 of Pex8 to be a major interaction site with the Pex2/Pex10/Pex12 RING finger complex. For reasons of space, labels P2, P5, P8, P10, and P12 are used as acronyms of Pex2, Pex5, Pex8, Pex10 and Pex12, respectively.
